# Global patterns of diversity and metabolism of microbial communities in deep-sea hydrothermal vent deposits

**DOI:** 10.1101/2022.08.08.503129

**Authors:** Zhichao Zhou, Emily St. John, Karthik Anantharaman, Anna-Louise Reysenbach

## Abstract

**Background:** When deep-sea hydrothermal fluids mix with cold oxygenated fluids, minerals precipitate out of solution and form hydrothermal deposits. These actively venting deep-sea hydrothermal vent deposits support a rich diversity of thermophilic microorganisms which are involved in a range of carbon, sulfur, nitrogen, and hydrogen metabolisms. Global patterns of thermophilic microbial diversity in deep-sea hydrothermal ecosystems have illustrated the strong connectivity between geological processes and microbial colonization, but little is known about the genomic diversity and physiological potential of these novel taxa. Here we explore this genomic diversity in 42 metagenomes from four deep-sea hydrothermal vent fields and a deep-sea volcano collected from 2004 to 2018, and document their potential implications in biogeochemical cycles.

**Results:** Our dataset represents 3,635 metagenome-assembled genomes encompassing 511 novel genera, with 395 Bacteria and 116 Archaea, providing many targets for cultivation of novel archaeal and bacterial families. Notably, 52% (206) of the novel bacterial genera and 72% (84) of the novel archaeal genera were found at the deep-sea Brothers volcano, many of which were endemic to the volcano. We report some of the first examples of medium to high-quality MAGs from phyla and families never previously identified, or poorly sampled, from deep-sea hydrothermal environments. We greatly expand the novel diversity of Thermoproteia, Patescibacteria (Candidate Phyla Radiation, CPR), and Chloroflexota found at deep-sea hydrothermal vents and identify a small sampling of two potentially novel phyla, designated JALSQH01 and JALWCF01. Metabolic pathway analysis of metagenomes provides insights into the prevalent carbon, nitrogen, sulfur and hydrogen metabolic processes across all sites, and illustrates sulfur and nitrogen metabolic ‘handoffs’ in community interactions. We confirm that Campylobacteria and Gammaproteobacteria occupy similar ecological guilds but their prevalence in a particular site is driven by shifts in the geochemical environment.

**Conclusion:** Our study of globally-distributed hydrothermal vent deposits provides a significant expansion of microbial genomic diversity associated with hydrothermal vent deposits and highlights the metabolic adaptation of taxonomic guilds. Collectively, our results illustrate the importance of comparative biodiversity studies in establishing patterns of shared phylogenetic diversity and physiological ecology, while providing many targets for enrichment and cultivation of novel and endemic taxa.

## Introduction

Actively venting deep-sea hydrothermal deposits at oceanic spreading centers and arc volcanoes support a high diversity of thermophilic microorganisms. Many of these microbes acquire metabolic energy from chemical disequilibria created by the mixing of reduced high temperature endmember hydrothermal fluids with cold oxygenated seawater. Community analysis of deposits using the 16S rRNA gene has revealed a rich diversity of novel archaeal and bacterial taxa [1–4] where the community composition is strongly influenced by the abundance of redox reactive species in high temperature vent fluids [e.g. 5–7]. The variations in the composition of endmember fluids, and in turn the microbial community composition at different vent fields, reflect the temperature and pressure of fluid-rock interaction, in addition to substrate composition and entrainment of magmatic volatiles. For example, along the Mid-Atlantic Ridge, methanogens are associated with deposits from H_2_-rich vents at Rainbow and are absent in H_2_-poor vents at Lucky Strike [3]. At the Eastern Lau Spreading Center (ELSC), similar to other back-arc basins, the hydrothermal fluids are generally quite variable depending on differences in inputs of acidic magmatic volatiles, contributions from the subducting slab and proximity of island arc volcanoes. Such geochemical differences are imprinted in the diversity of microbial communities [3, 4]. Similar complex community structure dynamics have also been recently reported for the communities of the submarine Brothers volcano on the Kermadec Arc [8].

While such global patterns of high temperature microbial diversity in deep-sea hydrothermal systems have demonstrated geological drivers of microbial colonization, little is known about the genomic diversity and physiological potential of the many reported novel taxa. While a few metagenomic studies of hydrothermal fluids and sediments have provided a much greater understanding of the functional potential of these communities [e.g. 7, 9–13], the metagenomic analysis of deposits has been limited to a small number of samples [e.g. 14–16]. One exception is the study of about 16 deep-sea hydrothermal deposits from Brothers volcano, which resulted in 701 medium and high quality metagenome-assembled genomes (MAGs) [8]. Further, this study demonstrated that there were functionally distinct high temperature communities associated with the volcano that could be explained through an understanding of the geological history and subsurface hydrologic regime of the volcano.

Here, we expand on the Brothers volcano study by exploring the genomic and functional diversity of hydrothermal deposits collected from deep-sea vents in the Pacific and Atlantic oceans. We greatly increase the number of novel high quality assembled genomes from deep-sea vents, many of which are endemic to vents and do not have any representatives in culture yet. We also show that known important biogeochemical cycles in hydrothermal ecosystems are accomplished by the coordination of several taxa as metabolic handoffs, where in some cases different taxa accomplish similar functions in different environments, potentially providing functional redundancy in fluctuating conditions.

## Results and Discussion

### Patterns of metagenomic diversity in deep-sea hydrothermal deposits

We sequenced 42 metagenomes from 40 samples (38 hydrothermal vent deposit samples and two diffuse flow fluids) collected at deep-sea hydrothermal vents and a deep-sea volcano. These represent one of the largest global collections of metagenomes from such samples (Fig. S1, S2). This study spans vent deposit collections from 2004 to 2018, from deep-sea hydrothermal vent fields in the north Atlantic (Mid-Atlantic Ridge, MAR), east and southwest Pacific (East Pacific Rise, EPR; Eastern Lau Spreading Center, ELSC), a sedimented hydrothermal system (Guaymas Basin, GB), and a deep-sea volcano (Brothers volcano, BV) (Table S1).

In this study, *de novo* assembly of sequencing data and subsequent genome binning and curation (see methods for details) resulted in 2,983 bacterial and 652 archaeal draft metagenome-assembled genomes (MAGs with ≥50% completeness, Table S2). Of these, ∼21% were >90% complete, with <5% contamination, and ∼36% contained a 16S rRNA gene fragment. The MAGs were initially characterized phylogenetically using the Genome Taxonomy Database Toolkit (GTDB-Tk) (Fig. 1, 2, 3, Data S1, S2, S3, S4, S5) [17]. MAGs that could not be assigned to a known genus by GTDB-Tk were assigned to new genera using AAI with the recommended cutoffs in Konstantinidis et al. [18] (Table S3A, B). Shared phyla between most of the hydrothermal deposits (excluding samples from the highly acidic Brothers volcano sites, and the diffuse flow fluids) included the Halobacteriota (e.g. Archaeoglobaceae), Methanobacteriota (e.g. Thermococcaceae), Thermoproteota (e.g. Acidilobaceae, Pyrodictiaceae), Acidobacteriota, Aquificota (e.g. Aquificaceae), Bacteroidota (e.g. Flavobacteriaceae), Campylobacterota (e.g. Sulfurimonadaceae, Nautiliaceae, Hippeaceae), Chloroflexota, Deinococcota (e.g. Marinithermaceae), Desulfobacterota (e.g. Dissulfuribacteraceae, Thermodesulfobacteriaceae), Proteobacteria (e.g. Alphaproteobacteria, Gammaproteobacteria), and the Patescibacteria (Table S4). Many of these phyla have only a few representatives in isolated cultures and point to the importance of combining enrichment cultivation strategies with metagenomic approaches to obtain additional insights into the physiological ecology of these core lineages.

**Fig. 1.**
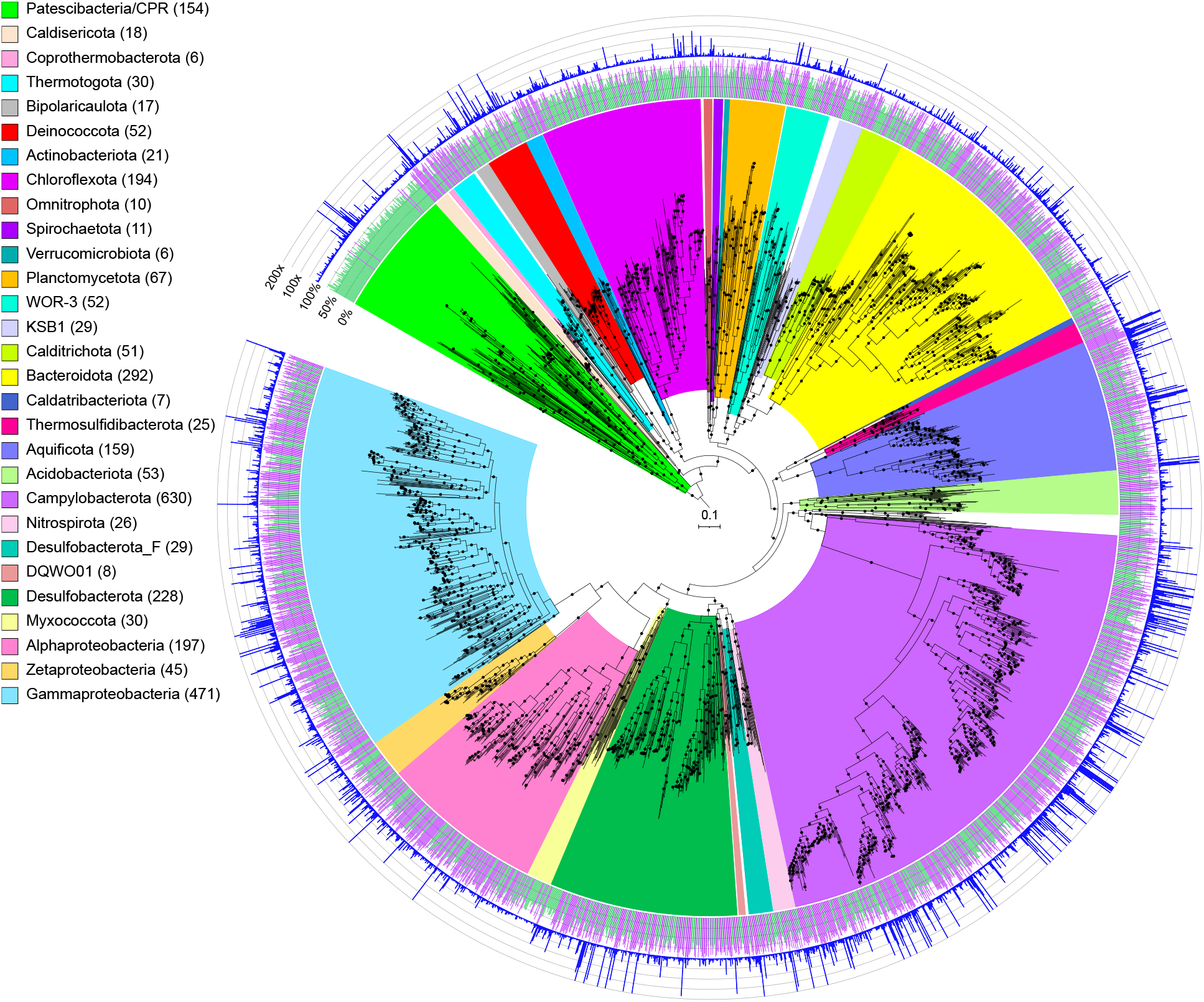
Maximum-likelihood phylogenomic tree of bacterial metagenome-assembled genomes, constructed using 120 bacterial marker genes in GTDB-Tk. Major taxonomic groups are highlighted, and the number of MAGs in each taxon is shown in parentheses. Bacterial lineages are shown at the phylum classification, except for the Proteobacteria which are split into their component classes. The inner ring displays estimated completion (green: 50-80%, purple: 80-100%), and the outer ring shows normalized read coverage up to 200x. The scale bar indicates 0.1 amino acid substitutions per site, and filled circles are shown for SH-like support values ≥80%. The tree was artificially rooted with the Patescibacteria using iTOL. The Newick format tree used to generate this figure is available in Data S4, and the formatted tree is available online at https://itol.embl.de/shared/alrlab.

**Fig. 2.**
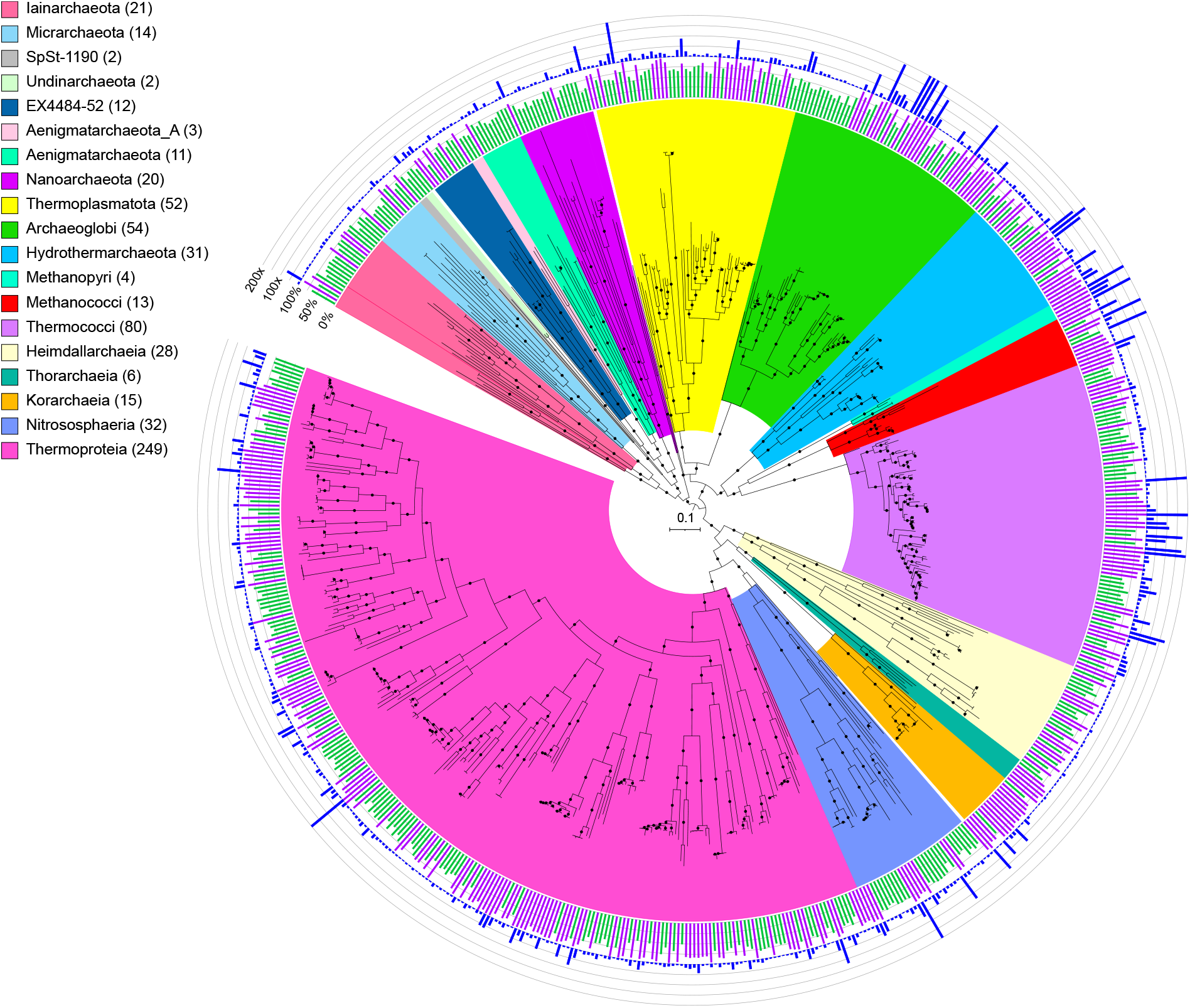
Maximum-likelihood phylogenomic reconstruction of deep-sea hydrothermal vent archaeal metagenome-assembled genomes generated in GTDB-Tk. The tree was generated with 122 archaeal marker genes. Taxa are shown at the phylum level, except for the Thermoproteota, Asgardarchaeota, Halobacteriota, and Methanobacteriota, shown at the class level. The number of MAGs in each highlighted taxon is shown in parentheses. Completion is shown on the inner ring (green: 50-80%, purple: 80-100%), while the outer ring displays normalized read coverage up to 200x. SH-like support values ≥80% are indicated with filled circles, and the scale bar represents 0.1 amino acid substitutions per site. The tree was artificially rooted with the Iainarchaeota, Micrarchaeota, SpSt-1190, Undinarchaeota, Nanohaloarchaeota, EX4484-52, Aenigmarchaeota, Aenigmarchaeota_A and Nanoarchaeota using iTOL. The tree used to create this figure is available in Newick format (Data S5), and the formatted tree is publicly available on iTOL at https://itol.embl.de/shared/alrlab.

**Fig. 3.**
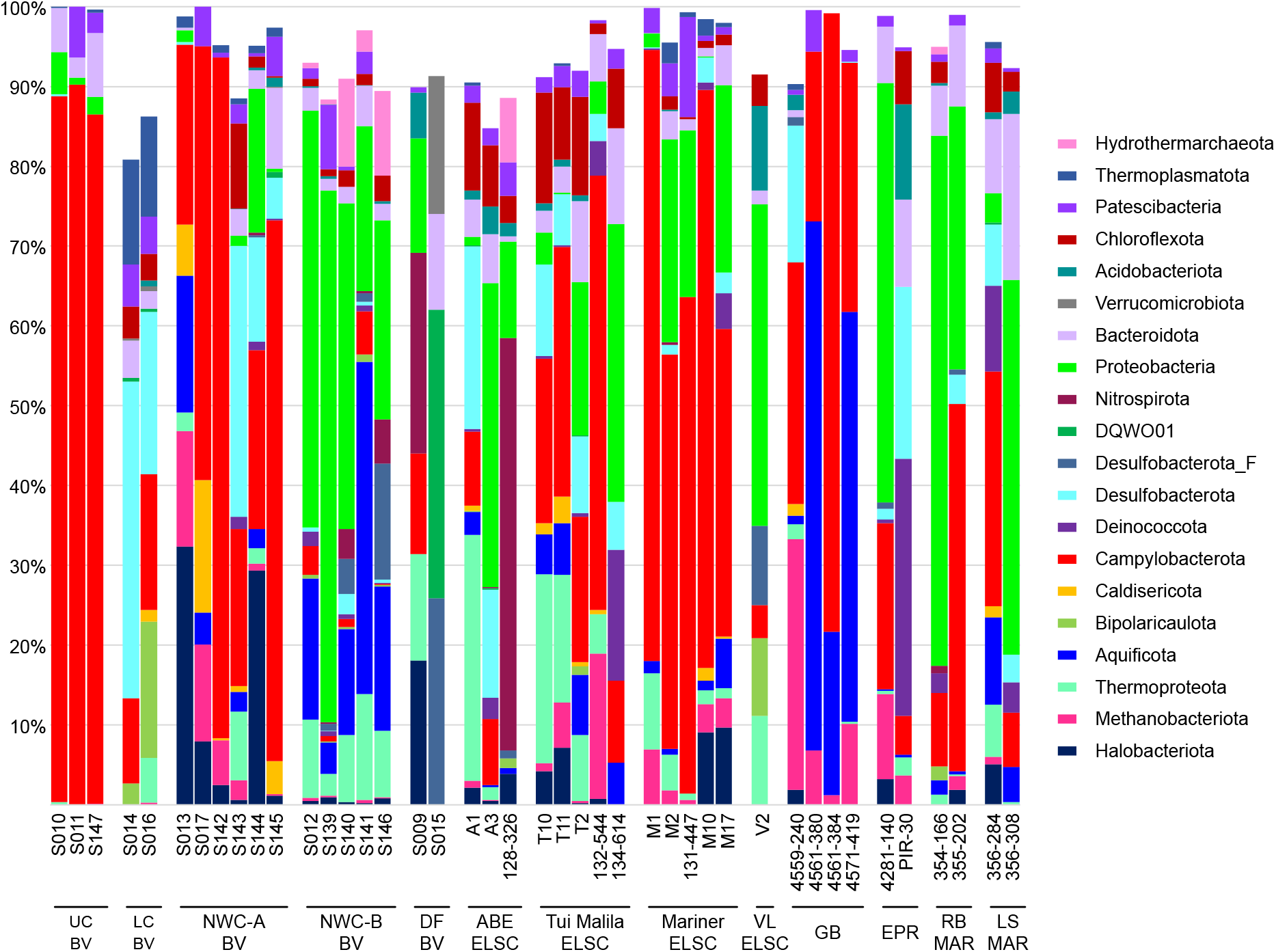
Relative abundance of MAG phyla, based on normalized read coverage. The phyla shown comprise ≥10% of the MAG relative abundance in at least one metagenomic assembly. Read coverage was normalized to 100M reads per sample, and coverage values for MAGs were summed and expressed as a percent. UC, Upper Cone; LC, Lower Cone, NWC-A, Northwest Caldera Wall A; NWC-B, Northwest Caldera Wall B and Upper Caldera Wall; DF, diffuse flow; VL, Vai Lili; RB, Rainbow; LS, Lucky Strike.

While shared taxa differed in relative abundance and distribution, observable differences in community structure between vent fields were somewhat limited in this study due to small sample numbers from some of the vent fields (two samples apiece from EPR; Rainbow, MAR; Lucky Strike, MAR), and the overall lower read depth of samples from these sites and a few other samples (Fig. S3). Therefore, obtaining statistically robust community structure patterns using MAG phylogenetic diversity for the entire dataset was not possible. However, Reysenbach et al. [8] did show that if metagenomic sequencing is deep, assembled MAG diversity tracks 16S rRNA amplicon diversity structure. Extrapolating to this study, the Brothers volcano MAG diversity patterns were retained and confirmed the amplicon observations from Reysenbach et al., 2020 [8] (Fig. S4), and in turn tracked the ELSC MAG community diversity (Fig. 4A, B). For example, sites at Brothers volcano that were hypothesized to have some magmatic inputs were predicted to be more similar in community structure to the sites along the ELSC with greater magmatic inputs, such as Mariner. Several of the samples from the more acidic Mariner vent field were more closely aligned in MAG diversity structure to those of the acidic solfataric Upper Cone sites at Brothers. The MAG data also demonstrated that the Guaymas samples were quite unique, which is not surprising, given that Guaymas Basin is a sediment-hosted system where the hydrothermal fluid geochemistry is quite different from other basalt- or andesitic-hosted hydrothermal systems (e.g., higher pH, high organics, high ammonia and methane) [19, 20].

**Fig. 4.**
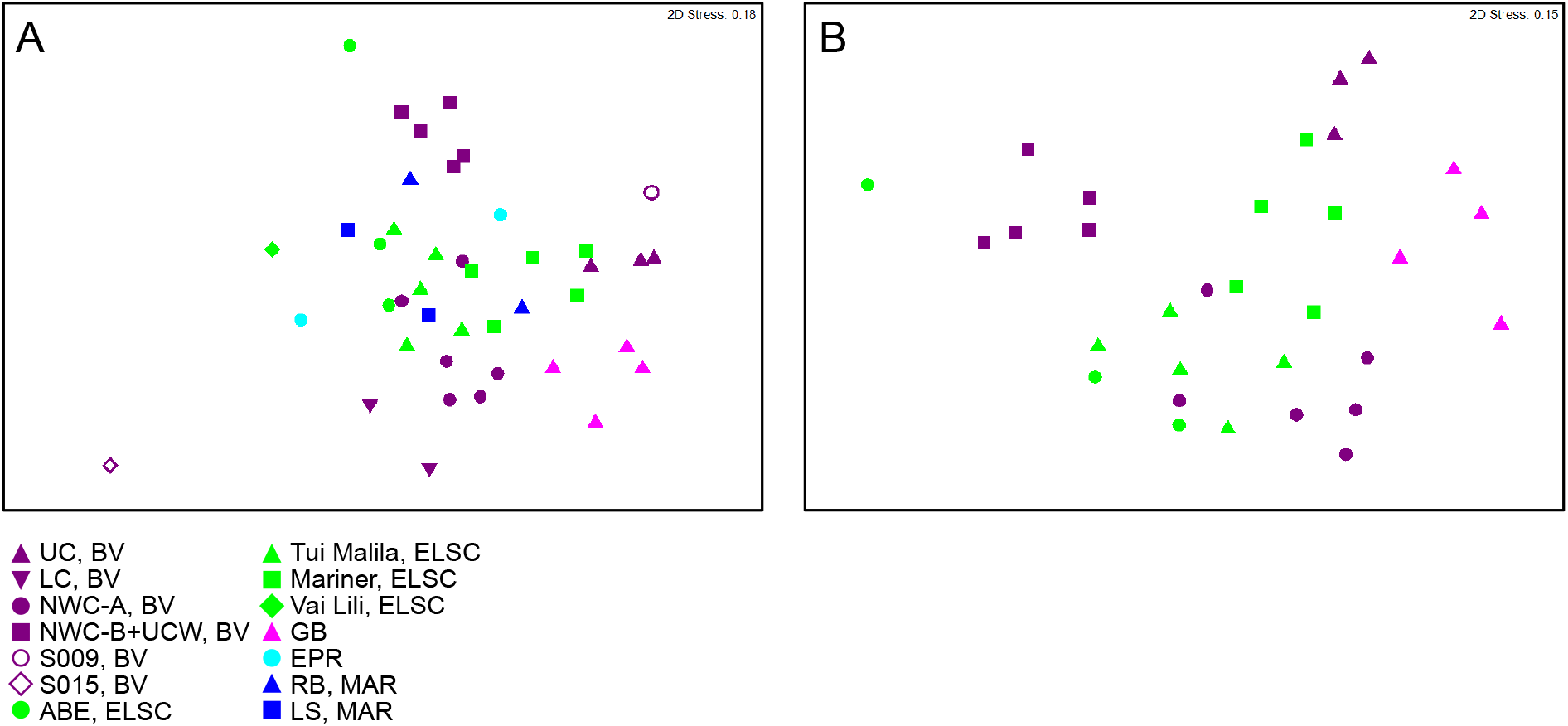
Non-metric multidimensional scaling (NMDS) plots showing taxonomic diversity of MAGs. Plots depict (A) all samples in this study and (B) a subset of the data, limited to locations with three or more samples. Plots were generated using Bray-Curtis matrices of the relative abundance of GTDB taxa, based on normalized read coverage of medium- and high-quality MAGs (Table S4; set to 100M reads and expressed as a percentage of MAG read coverage per sample). Points that are closer together in the plots represent a higher degree of similarity.

Our dataset greatly broadens genomic diversity from deep-sea vents, by representing 511 novel genera, comprising 395 Bacteria and 116 Archaea. Notably, 52% (206) of the novel bacterial genera and 72% (84) of the novel archaeal genera were found at Brothers volcano. Furthermore, 25% (99) of the novel bacterial genera and 47% (54) of the novel archaeal genera were unique to the Brothers volcano samples, which further supports the understanding that this environment is a hotbed for novel microbial biodiversity, reflected in the volcano’s complex subsurface geology [8].

While many of these novel archaeal and bacterial genera were previously reported from Brothers volcano [8], we report them again here in the context of the new assemblies (1000 bp contig cutoff, used for Brothers volcano samples and ELSC 2015 samples) and iterative DAS Tool binning used for all our metagenomes. Our data support that of Reysenbach et al., [8], which used MetaBAT for assemblies (2000 bp contig cutoff) of the Brothers volcano metagenomes. Namely, we recovered approximately 202 novel bacterial genera and 83 new archaeal genera from Brothers volcano communities in Reysenbach et al. [8], well within the range detected in this analysis (viz. 206 and 84, respectively). In this study, using a lower contig cutoff allowed for the recovery of a much higher number of MAGs, but many are of lower quality with higher contig counts. For example, MAGs recovered in the Reysenbach et al. [8] study had an average of 254 contigs per MAG, with ∼19% (135) of MAGs comprising 100 contigs or less. In contrast, only 7% (258) of MAGs in this current study had 100 contigs or less, and the average number of contigs per MAG was 511 (Table S2). However, using the iterative binning approach provided advantages when resolving lineages of high microdiversity, such as in the Nautiliales, with the caveat of creating some MAGs with large collections of erroneous contigs that were poorly detected by CheckM, as they had very few associated marker genes (e.g. MAGs 4571-419_metabat1_scaf2bin.008, M10_maxbin2_scaf2bin.065; Fig. S5). This points to the importance of carefully choosing assembly parameters depending on the ultimate goal of whether quality over quantity of MAGs is preferred for analyses of ecological patterns. Our data demonstrate, however, that overall patterns of MAG diversity are retained regardless of assembly techniques and parameters (Fig. S4).

Furthermore, here we document some of the first examples of medium to high quality MAGs from phyla and classes never previously identified, or poorly sampled, from deep-sea hydrothermal environments. These include Thermoproteia, Patescibacteria (formerly Candidate Phyla Radiation, CPR), Chloroflexota, and a few MAGs representing two putative new bacterial phyla, JALSQH01 (3 MAGs) and JALWCF01 (13 MAGs) (Supplementary Discussion, Fig. S6, Table S5). For example, with 249 MAGs belonging to the Thermoproteia (Table S2, Fig. S7), we have significantly expanded the known diversity and genomes from this phylum. The importance of this group at deep-sea vents was first recognized through 16S rRNA amplicon studies, where the depth of sequencing highlighted that much of this novel thermophilic diversity had been overlooked [e.g. 3, 4]. Furthermore, it is now recognized that many members of this group have several introns in the 16S rRNA gene, which explains why they were missed in original clone library assessments and may be underestimated in amplicon sequencing [21–24]. For example, 24 MAGs were related to a recently described genus of the Thermoproteia, *Zestosphaera* (GTDB family NBVN01) [24]. This genus was first isolated from a hot spring in New Zealand but is clearly a common member of many deep-sea vent sites. Further, the discovery of a 16S rRNA gene related to *Caldisphaera* at deep-sea vents [25], previously only detected in terrestrial acidic solfataras, led to the isolation of related Thermoplasmata – *Aciduliprofundum boonei* – but the *Caldisphaera* escaped cultivation. Here we report several high-quality MAGs related to this genus (M2_metabat2_scaf2bin.319, 131-447_metabat1_scaf2bin.050, M1_metabat1_scaf2bin.025, S016_metabat2_scaf2bin.003). Additionally, we also recovered a genome from the Gearchaeales (S146_metabat1_scaf2bin.098), first discovered in iron-rich acidic mats in Yellowstone National Park [26], and members of the poorly sampled Ignicoccaceae, Ignisphaeraceae, and Thermofilaceae. While we identified several genomes from recently discovered archaeal lineages including the Micrarchaeota, Iainarchaeota and Asgardarchaeota, we also recovered 15 MAGs belonging to the Korarchaeia, 14 of which comprise two putative novel genera, and one which is closely related to a MAG previously recovered from sediment in Guaymas Basin (Genbank accession DRBY00000000.1) [27, 28]. Additionally, we recovered four MAGs from the Caldarchaeales that span two novel genera, one of which was recently was proposed as *Candidatus* Benthortus lauensis [29] using a MAG generated from a previous assembly of the T2 metagenome (T2_175; Genbank accession JAHSRM000000000.1). MAGs belonging to this genus were identified at both Tui Malila, ELSC, and Brothers volcano (T2_metabat2_scaf2bin.284, S140_maxbin2_scaf2bin.281, S141_maxbin2_scaf2bin.262) with the Tui Malila MAG nearly identical (99.7% AAI similarity) to the described *Cand*. B. lauensis T2_175 MAG.

While within the Bacteria, the Gammaproteobacteria and Campylobacterota were by far the most highly represented bacterial genomes, there were other lineages for which we have very little if any data or cultures from deep-sea hydrothermal systems (Fig. 3, Fig. S7). Two such groups are the Patescibacteria and Chloroflexota, with 154 and 194 MAGs respectively.

### Patescibacteria and Chloroflexota are diverse and abundant members of deep-sea hydrothermal vent deposits

The Patescibacteria/Candidate Phyla Radiation (CPR) encompasses a phylogenetically diverse branch within the bacterial tree of life that is poorly understood and rarely documented in deep-sea hydrothermal systems. Originally, the CPR was proposed to include several phylum-level lineages [30], but the entire group was later reclassified by GTDB as a single phylum, Patescibacteria [31]. Members of the Patescibacteria have been well-characterized in terrestrial soils, sediments, and groundwater [32–37], and in the mammalian oral cavity [38–40]. Several 16S rRNA gene and metagenomic studies have also identified members of the Patescibacteria from deep-sea vents, including EPR, MAR, ELSC, and Guaymas Basin [3, 4, 12, 15, 41–43], from Suiyo Seamount [44], and the Santorini submarine volcano [45], further supporting the widespread distribution of this metabolically diverse phylum.

Our study adds 56 novel genera based on AAI and GTDB classifications to the Patescibacteria phylum. These include large clades within the Gracilibacteria (10 new genera), representatives within the Microgenomatia (9 novel genera), Dojkabacteria (10 new genera), and several clades in the Paceibacteria (13 new genera) (Fig. 5A, B, Fig. S8). The Gracilibacteria and Paceibacteria were overall the most prevalent lineages of Patescibacteria in the samples but had contrasting distributions across vents (Fig. 5B). In general, when the Gracilibacteria were prevalent, the Paceibacteria appeared to be a minor component or not present, and vice versa. In particular, the Gracilibacteria MAGs were often associated with the acidic sites such as the Upper Cone at Brothers volcano (S011, S147), and the Mariner vent fields, and in the early colonization experiment from Guaymas Basin (Supplementary Discussion). This may suggest that Gracilibacteria function as early colonizers and are associated with turbulent ephemeral environments as observed previously in oil seeps [46]. Continued investigation into the ecology, evolution, and host association patterns of these groups, however, may shed more light on these distribution differences.

**Fig. 5.**
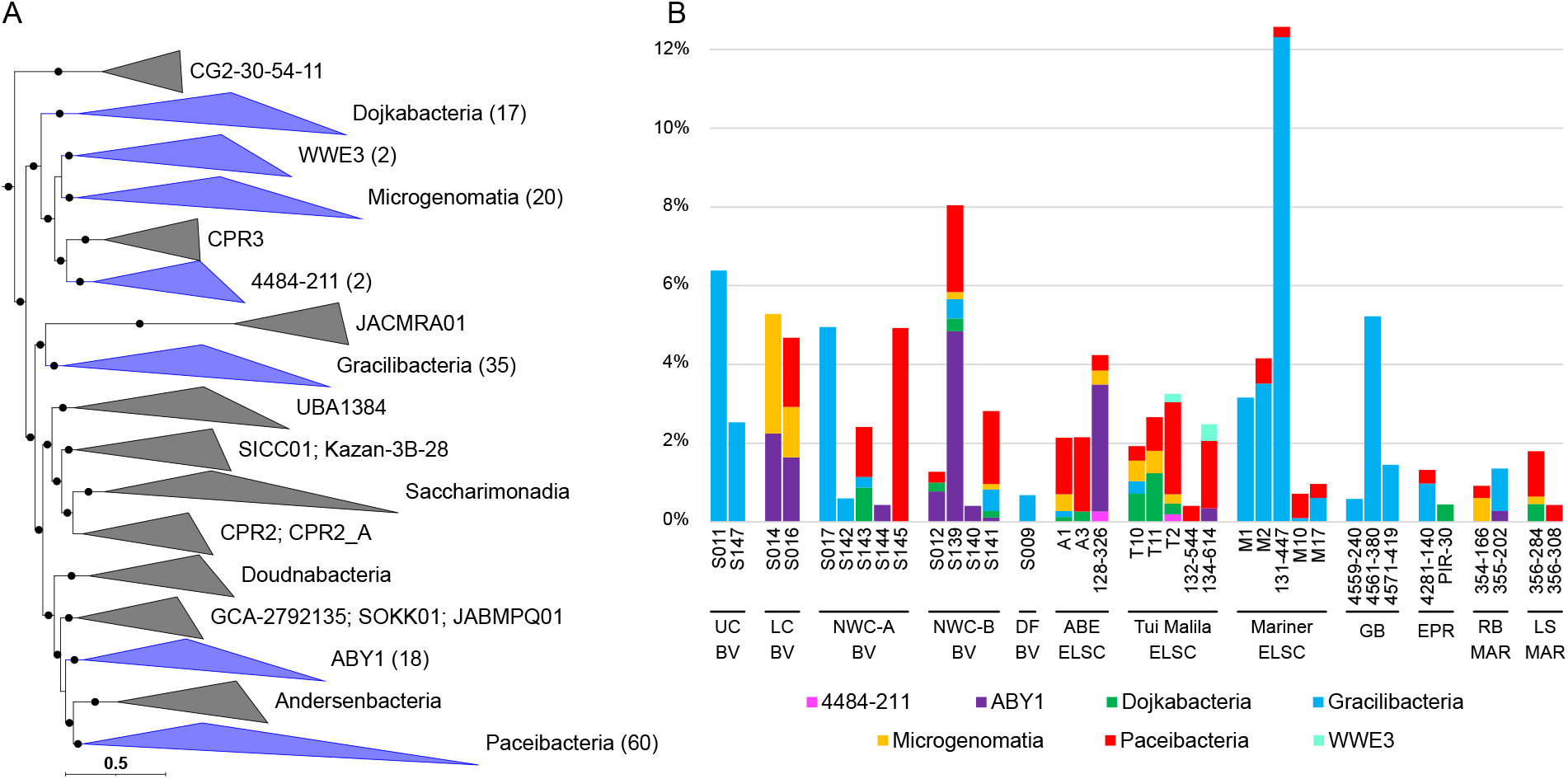
Phylogenomic placement and relative abundance of Patescibacteria MAGs, displayed at the class rank. (A) Blue clades in the maximum-likelihood phylogenomic tree contain MAGs from this study, with the number of MAGs shown in parentheses. The scale bar shows 0.5 substitutions per amino acid, and filled circles indicate SH-like support (≥80%). (B) Relative abundance of Patescibacteria MAGs was calculated using normalized read coverage for MAGs in each assembly (set to 100M reads and expressed as a percentage of MAG read coverage per sample).

Consistent with previous studies [30, 34], many of the recovered Patescibacteria MAGs had very small genomes (often ∼1MB or smaller; Table S2) with highly reduced metabolic potential, often lacking detectable genes for synthesis of fatty acids, nucleotides, and most amino acids (Table S6). Gene patterns also suggested that many of the organisms are obligate anaerobes, lacking aerobic respiration, and that they likely form symbiotic or parasitic associations with other microbes, as has been shown for Patescibacteria cultivated thus far from the Absconditabacterales and Saccharibacteria [39, 40, 47, 48].

We recovered several MAGs from Mariner, Guaymas Basin, and Brothers volcano that were related to the parasitic *Cand*. Vampirococcus lugosii [47] and *Cand*. Absconditicoccus praedator [48]. In order to explore if our MAGs had any hints of a parasitic lifestyle, we searched for some of the large putative cell-surface proteins identified in the genomes of *Cand*. V. lugosii [47] and *Cand*. A. praedator [48]. Using a local BlastP of nine of the longest genes found in *Cand*. V. lugosii, we recovered high-confidence homologs (E-value=0) for alpha-2 macroglobulin genes in several MAGs from the Abscontitabacterales (based on search of *Cand*. V. lugosii protein MBS8121711.1), which may be involved in protecting parasites against host defense proteases [47]. We also recovered homologs for PKD-repeat containing proteins (MBS8122536.1; E-value=0), which are likely involved in protein-protein interactions [47]. Previous analysis of *Cand*. V. lugosii found these giant proteins are likely membrane-localized, suggesting they may potentially play a role in host/symbiont interactions. Additionally, we identified these long proteins from *Cand*. V. lugosii elsewhere in the Gracilibacteria MAGs. For example, putative homologs of the PKD repeat containing protein (MBS8122536.1), a hypothetical protein (MBS8121701.1), and the alpha-2 macroglobulin (MBS8121711.1) were identified in multiple other orders of the class Gracilibacteria (E-value ≤1E-25). The alpha-2 macroglobulin was also identified in the very distantly related Paceibacteria, and a single putative homolog of the alpha-2 macroglobulin was found in a MAG belonging to the class WWE3 (134-614_metabat1_scaf2bin.084; E-value ≤1E-24).

While the Patescibacteria likely rely on symbiotic or parasitic relationships, members of the Chloroflexota phylum are diverse and metabolically flexible organisms, capable of thriving in a wide variety of geochemical niches. Chloroflexota are abundant and widely distributed in a variety of environments, including terrestrial soils, sediments and groundwater, freshwater, pelagic oceans, and the marine subseafloor and sediments [49–55], and hydrothermal settings such as Guaymas Basin [11] and Brothers submarine volcano [8]. Genomic evidence suggests that Chloroflexota are associated with important metabolisms in the carbon cycle, including fermentation, carbon fixation, acetogenesis and the utilization of sugars, polymers, fatty acids, organic acids and other organic carbon compounds [50, 51, 54].

Here we add to the growing evidence that the Chloroflexota are diverse and metabolically versatile members of deep-sea hydrothermal vent communities. We recovered a total of 194 Chloroflexota MAGs spanning 12 orders (GTDB taxonomy), which included 22 novel genera. Of these novel genera, 14 were identified at Brothers volcano and 6 were unique to the Brothers volcano samples (Table S3A). Based on read coverage, Chloroflexota MAGs were in high relative abundance (≥7%) in several samples from the ELSC, namely, from Tui Malila and ABE, and in one NW Caldera Wall sample from Brothers volcano (Table S4). To further explore the metabolic potential of Chloroflexota in hydrothermal vent communities, we focused our analyses on high-completeness MAGs (≥80% completeness, *n*=58) distributed in 6 orders: Caldilineales, Promineofilales, Anaerolineales, Ardenticatenales, B4-G1, and SBR1031 (Fig. 6, Table S7A).

**Fig. 6.**
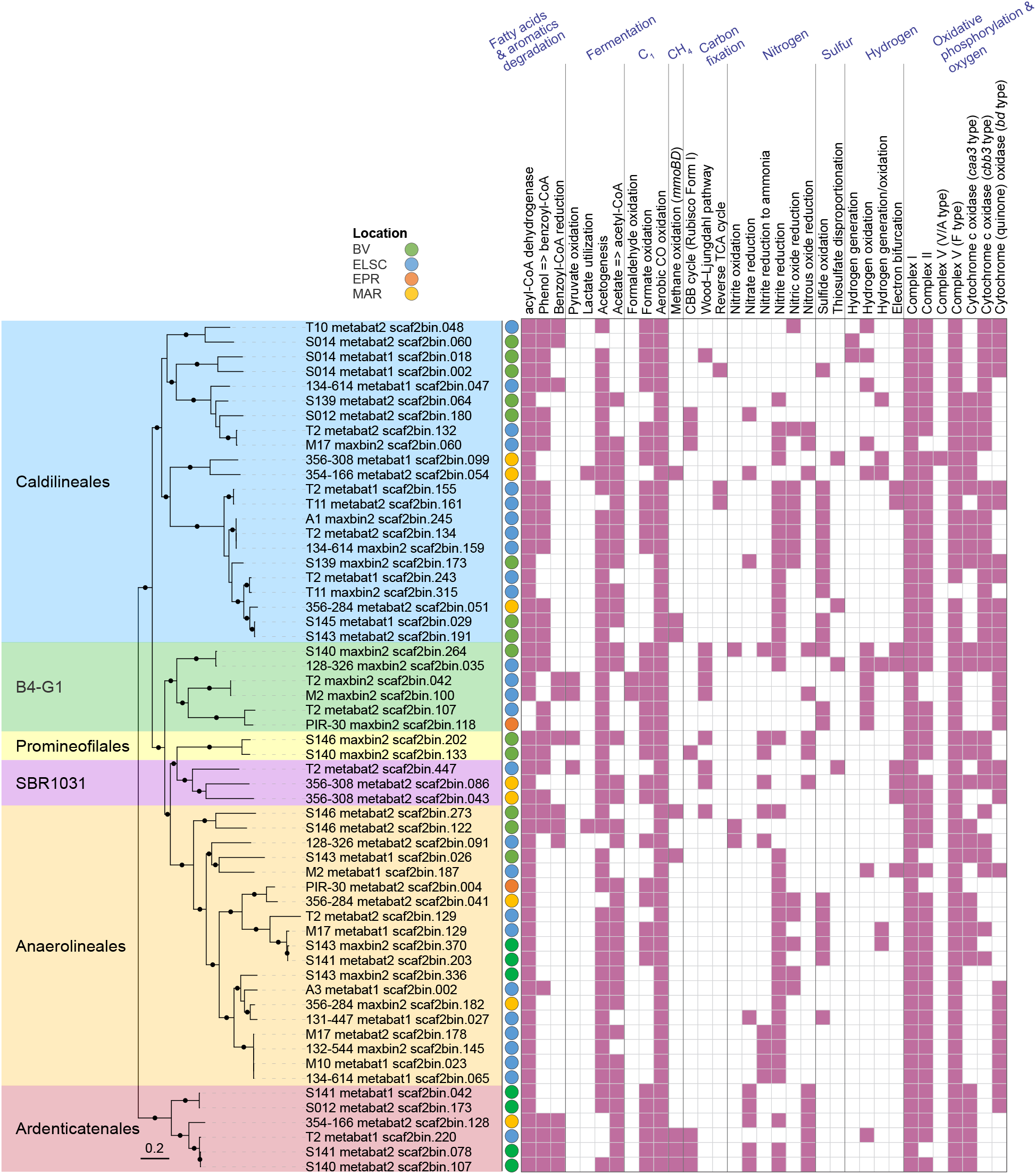
Phylogenetic tree of 58 high-completeness (≥80%) Chloroflexota MAGs with predicted functional capabilities. Nodes with ultrafast bootstrap support values ≥90% are shown with filled circles, and the scale bar shows 0.2 substitutions per site. One genome from the GTDB r202 database (GTDB accession GB_GCA_007123655.1) was used to re-root the tree.

The majority (≥75%) of the high quality Chloroflexota MAGs encoded marker genes involved in several processes previously associated with the Chloroflexota (Table S7B), including fatty acid degradation [50, 55], formate oxidation [56], aerobic CO oxidation [57] and selenate reduction [53]. Except for the Anaerolineales, over 66% of the MAGs in the other five orders had the capacity for degradation of aromatic compounds, as previously reported for Chloroflexi from the marine subsurface [51]. While some MAGs had the potential for substrate-level phosphorylation through acetate formation, most of the MAGs contained pathways for oxidative phosphorylation and oxygen metabolism [50, 51]. The Wood–Ljungdahl pathway, the CBB cycle based on a Form I Rubisco, and the reverse TCA cycle were detected in some of the MAGs [50, 51]. Soluble methane monooxygenase genes, a metabolic potential recently also detected in a Chloroflexota MAG from the arctic [58], were identified in a total of eight of our MAGs from the orders Caldilineales, Anaerolineales, and Ardenticatenales.

Although the primary metabolic potential of the hydrothermal vent-associated Chloroflexota was in carbon cycling, we did, however, observe minor evidence for their roles in nitrogen and sulfur cycling (Fig. 6, Table S7). About 22% of the MAGs (with ≥80% completeness) encoded capacities for sulfide oxidation, as previously reported for members of this group, e.g. *Chloroflexus* spp. [59, 60]. The potential to disproportionate thiosulfate was also observed in a few MAGs. Further, thermophilic Chloroflexi grown in an enrichment culture from Yellowstone National Park were shown to oxidize nitrite. A few of our MAGs encoded genes involved in nitrite oxidation [61], while a larger proportion of the MAGs encoded genes for nitrite or nitric oxide reduction. None of the MAGs encoded complete pathways for entire sulfur oxidation or denitrification, suggesting that Chloroflexota in these environments may be associated with metabolic handoffs involving other community members (see below).

### Metabolic and functional diversity in deep-sea hydrothermal vent deposits

In order to explore the metabolic and functional diversity associated with our MAGs, we utilized functional assignment results in tandem with the corresponding MAG relative abundance (Table S8). In general, genes involved in carbon, nitrogen, sulfur, and hydrogen metabolism were prevalent and shared across all hydrothermal systems in this study (Fig. 7, 8). While heterotrophy, autotrophy and mixotrophy potential were identified in all samples, 47.1% of the MAGs (by count) exhibited potential for carbon fixation. Marker genes associated with five different carbon fixation pathways were identified in the MAGs, namely, the Calvin-Benson-Bassham (CBB) cycle (form I or form II Rubisco), the 3-hydroxypropionate/4-hydroxybutyrate cycle, the dicarboxylate/4-hydroxybutyrate cycle, the reverse TCA cycle, and the Wood– Ljungdahl pathway (Fig. 7, 8). Marker gene presence also suggested the potential for widespread heterotrophic metabolism of peptides, polysaccharides, nucleotides and lipids, and fermentation via acetogenesis (Fig. 7, 8).

**Fig. 7.**
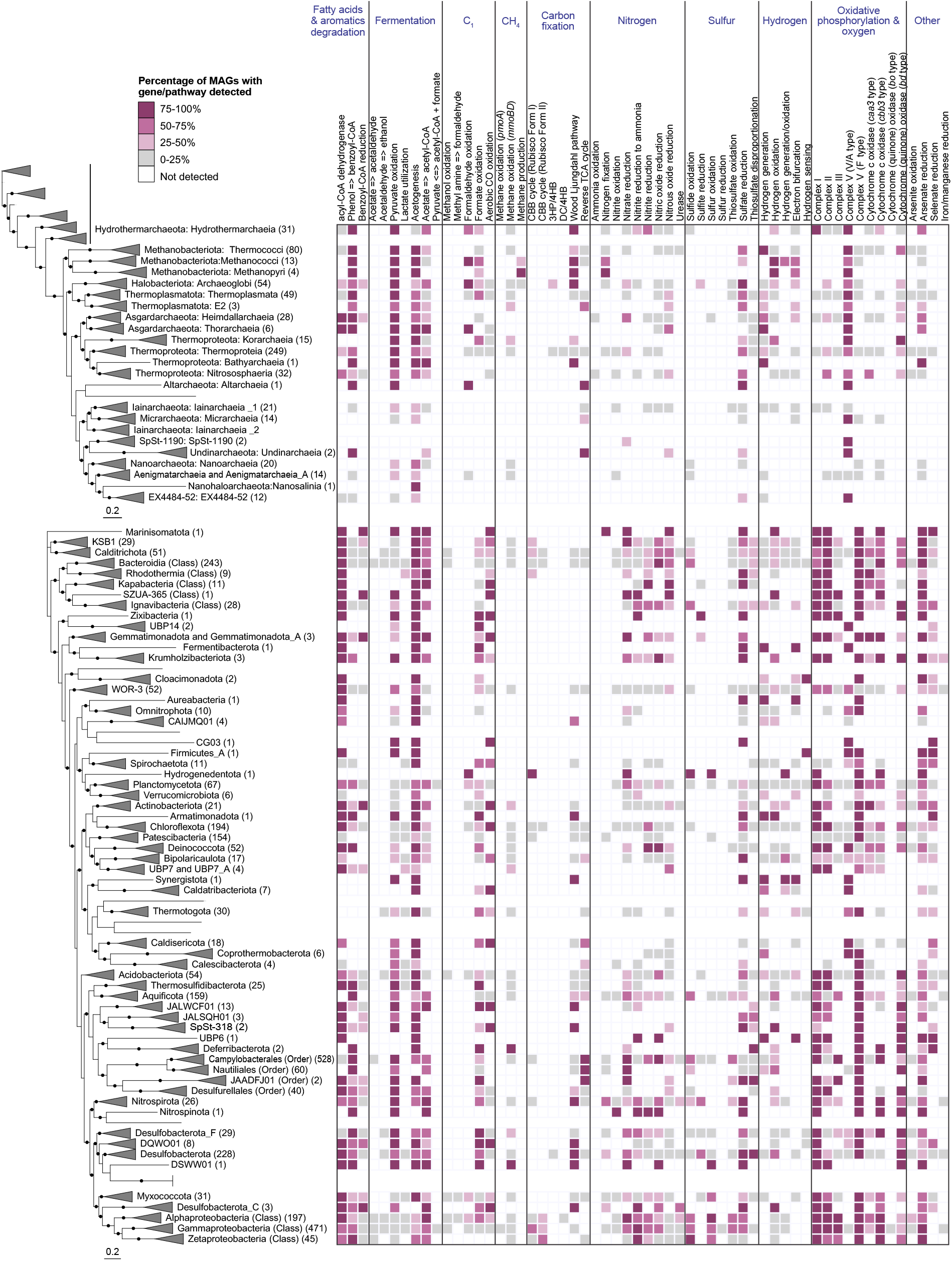
Core metabolic gene presence across phylogenetic clusters in deep-sea hydrothermal vent deposits. The number of MAGs in each clade is shown in parentheses, and MAGs belonging to unclassified lineages or falling outside their corresponding phylogenetic cluster due to unstable tree topology are shown without names. In instances where a phylum was not recovered as a monophyletic lineage within the tree (e.g. Iainarchaeia), MAG count and gene distribution for the entire phylum is only shown on one of the branches. Unless otherwise indicated, archaeal clades are shown at the class level, while bacterial clades are shown at the phylum level. Nodes with ultrafast bootstrap support ≥90% are shown with filled circles, and scale bars indicating 0.2 amino acid substitutions per site are provided for both archaeal and bacterial trees. Detailed metabolic gene presence information can be found in Table S9.

**Fig. 8.**
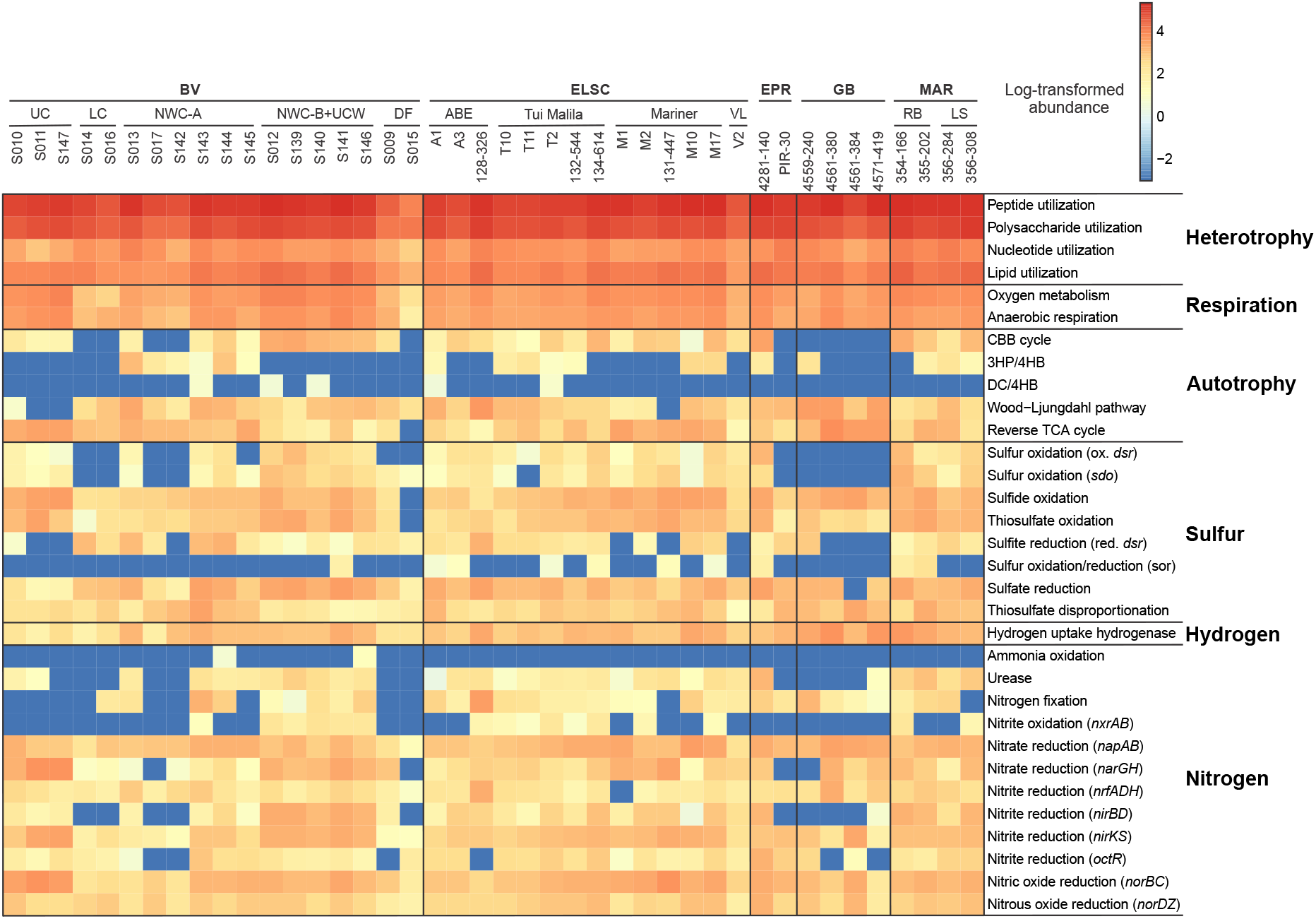
Heatmap displaying the metabolic potential for each metagenome. Within each metagenomic dataset, functional abundance values were calculated as described in the methods. Functional abundances were then log-transformed, with abundance values equal to zero replaced by 10^-3^ to avoid negative infinite values.

Genes involved in nitrogen fixation, denitrification, and nitrite oxidation were identified across the different hydrothermal sites, yet the potential for anaerobic or aerobic ammonia oxidation was rarely detected (Fig. 8). The absence of ammonia oxidation is not totally surprising, since ammonia is in very low to undetectable concentrations in deep-sea hydrothermal fluids, with the exception of sediment-hosted hydrothermal areas like at Guaymas Basin [19, 20]. In these sedimented hydrothermal systems, aerobic and anaerobic ammonia oxidation are key processes within the sediments and hydrothermal plumes [62–65], but they may not be as important in the hydrothermal deposits. Our data also expands the importance of nitrogen fixation from the first detection at deep-sea vents in *Methanocaldococcus* [66] to a greater diversity of hydrothermal Bacteria and Archaea.

Given the importance of sulfur cycling in deep-sea hydrothermal systems [67–69], it is not surprising that genes associated with elemental sulfur, sulfide, and thiosulfate oxidation, sulfate reduction, and thiosulfate disproportionation were widely distributed in MAGs from different hydrothermal samples and were associated with diverse taxonomic guilds (Fig. 7, 8). Based on metabolic gene distribution statistics (Table S9), the potential for sulfur oxidation was identified in 16% of the MAGs (577), primarily in members of the Alphaproteobacteria and Gammaproteobacteria. Genes associated with sulfide oxidation were identified in 34% of the MAGs (1216), including members of the Bacteroidia, Campylobacteria, Alphaproteobacteria, and Gammaproteobacteria. Thiosulfate oxidation genes were detected in 23% of the MAGs (836), largely comprised of the Campylobacteria, Alphaproteobacteria, and Gammaproteobacteria, while 14% of the MAGs (522) encoded genes for thiosulfate disproportionation, including the classes Bacteroidia, Campylobacteria and the phylum Desulfobacterota. The potential for dissimilatory sulfite reduction was identified in 6% of the MAGs (220) distributed across ten bacterial and archaeal phyla, namely Halobacteriota (class Archaeoglobi), Bacteroidota (class Kapabacteria), Campylobacterota (class Campylobacterales), Zixibacteria, Gemmatimonadota, Acidobacteriota, Nitrospirota, Desulfobacterota, Desulfobacterota_F and Myxococcota.

Hydrogen is highly variable in hydrothermal fluids, with some of the highest concentrations in geothermal systems hosted by ultramafic rocks, such as the Rainbow hydrothermal vent field [3], or in sediment-hosted regions like Guaymas basin [70]. In these systems, methanogens and sulfate reducers are prevalent hydrogen consumers [3, 71–74], although a wide variety of other heterotrophs and autotrophs can also derive energy from hydrogen oxidation [72]. Hydrogenase enzymes are responsible for mediating hydrogen oxidation in microbial populations but are also involved in a variety of other functions, including hydrogen evolution, electron bifurcation, and hydrogen sensing [75]. Approximately 27% of the MAGs in this study (974) encoded for at least one hydrogenase gene, and the MAGs were predominantly associated with the classes Campylobacteria, Bacteroidia, Gammaproteobacteria, and the phylum Desulfobacterota (Fig. 7, 8, Table S9). In several cases (132 MAGs), hydrogenase genes co-occurred with genes involved in the oxidation of reduced sulfur species (sulfide, elemental sulfur, sulfite or thiosulfate). This is not surprising, given that the capability to oxidize both sulfur and hydrogen has been shown in multiple isolates, including members of the Campylobacteria [76–78] and Aquificae [e.g. 79, 80].

### Metabolic handoffs are a central feature of community interactions in deep-sea hydrothermal vent deposits

The microbial communities at deep-sea hydrothermal vents are shaped by a wide variety of complex interactions, including symbiosis, syntrophy, commensalism, cross-feeding, and metabolic handoffs [11, 12, 81–83]. While many of the MAGs encode genes associated with different biogeochemical cycles, as expected, the genes for a complex functional pathway were not localized in a single MAG, but instead distributed across several MAGs. This is likened to ‘metabolic handoffs’ where the interaction between different organisms produces pathway intermediates, enabling community members to perform downstream reactions in the metabolic pathway. For example, metagenomic analysis of a subsurface aquifer environment suggested that metabolic handoffs are commonly utilized in key biogeochemical pathways such as sulfide oxidation and denitrification [37]. Genes for sulfide oxidation were identified in all the deep-sea hydrothermal vent sites in this study, but few MAGs encoded genes for the entire three-step pathway. A much larger proportion of the MAGs, however, contained genes for a single step in sulfur oxidation (Fig. 9), consistent with a metabolic handoff scenario. Similar patterns were also observed for sulfate reduction and denitrification (Fig. 9). Additionally, the genes for individual steps in sulfide oxidation were often found coupled with at least one gene from the denitrification pathway, which may increase the thermodynamic favorability of both pathways (Li et al., 2018). Additionally, one or more denitrification genes co-occurred with sulfide oxidation genes in 1113 MAGs, with elemental sulfur oxidation genes in 485 MAGs and with sulfite oxidation genes in 1025 MAGs (Table S9).

**Fig. 9.**
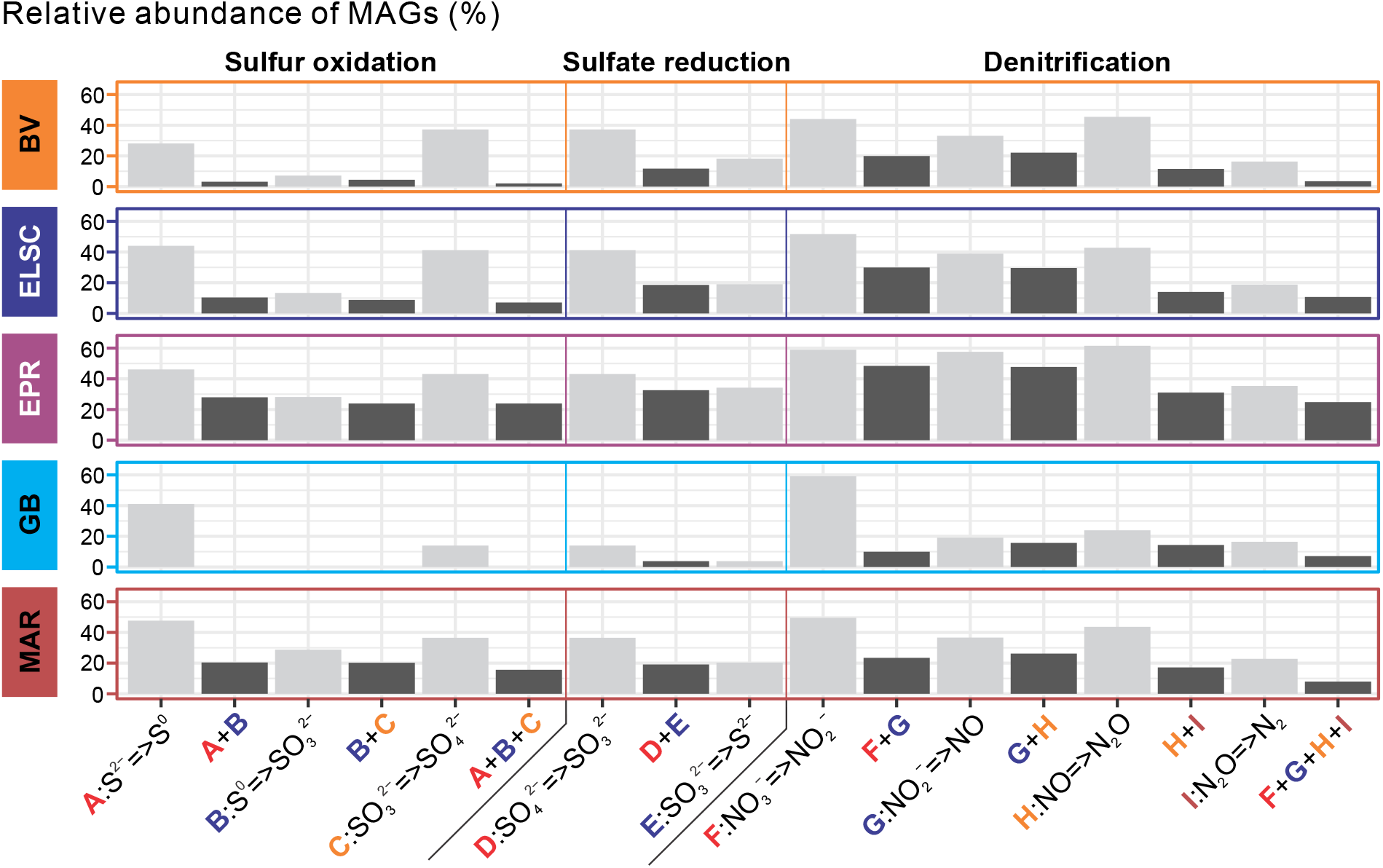
Bar plots showing the sequential steps of sulfur oxidation, denitrification and sulfate reduction. Bar height indicates the percent relative abundance of MAGs in each metagenome with genes for a particular function(s), averaged across hydrothermal vent sites.

### Conserved microbial functions are mediated by different taxa at different hydrothermal vent systems

Previous analyses of deep-sea hydrothermal environments and global oceans have pointed to widespread functional redundancy in microbial communities [8, 12, 84, 85], with similar metabolic potential identified across taxonomically diverse samples. For example, a study of Guaymas Basin metagenome-assembled genomes suggested that many functional genes could be identified across multiple distinct taxa [12]. In our study, members of the Campylobacteria and Gammaproteobacteria were present in almost all samples, yet showed contrasting patterns of abundance (Fig. 10). These lineages can perform several of the same functional processes including oxidation of reduced sulfur species [86], denitrification [87–89], and carbon fixation [90–93]. This can be partially explained by ecophysiological and growth differences between the groups, which are selected for by the different geochemical profiles at the various vent sites. For example, studies have suggested that Campylobacteria tend to favor higher sulfide conditions but have a broader range of oxygen tolerance than the Gammaproteobacteria, while Gammaproteobacteria tend to inhabit a narrower range of higher oxygen and lower sulfide [16, 86, 90, 94]. It is therefore not surprising that the Campylobacteria were more prevalent at several of the acidic and more turbulent sites, such as at the Upper Cone, Brothers volcano and in early colonized samples from a thermocouple array at Guaymas Basin (Table S4, Supplementary Discussion). Patwardhan et al. [95] also showed that Campylobacteria were early colonizers of shallow marine vents followed by Gammaproteobacteria, and their differential colonization could be linked to sulfide, oxygen, and temporal differences.

**Fig. 10.**
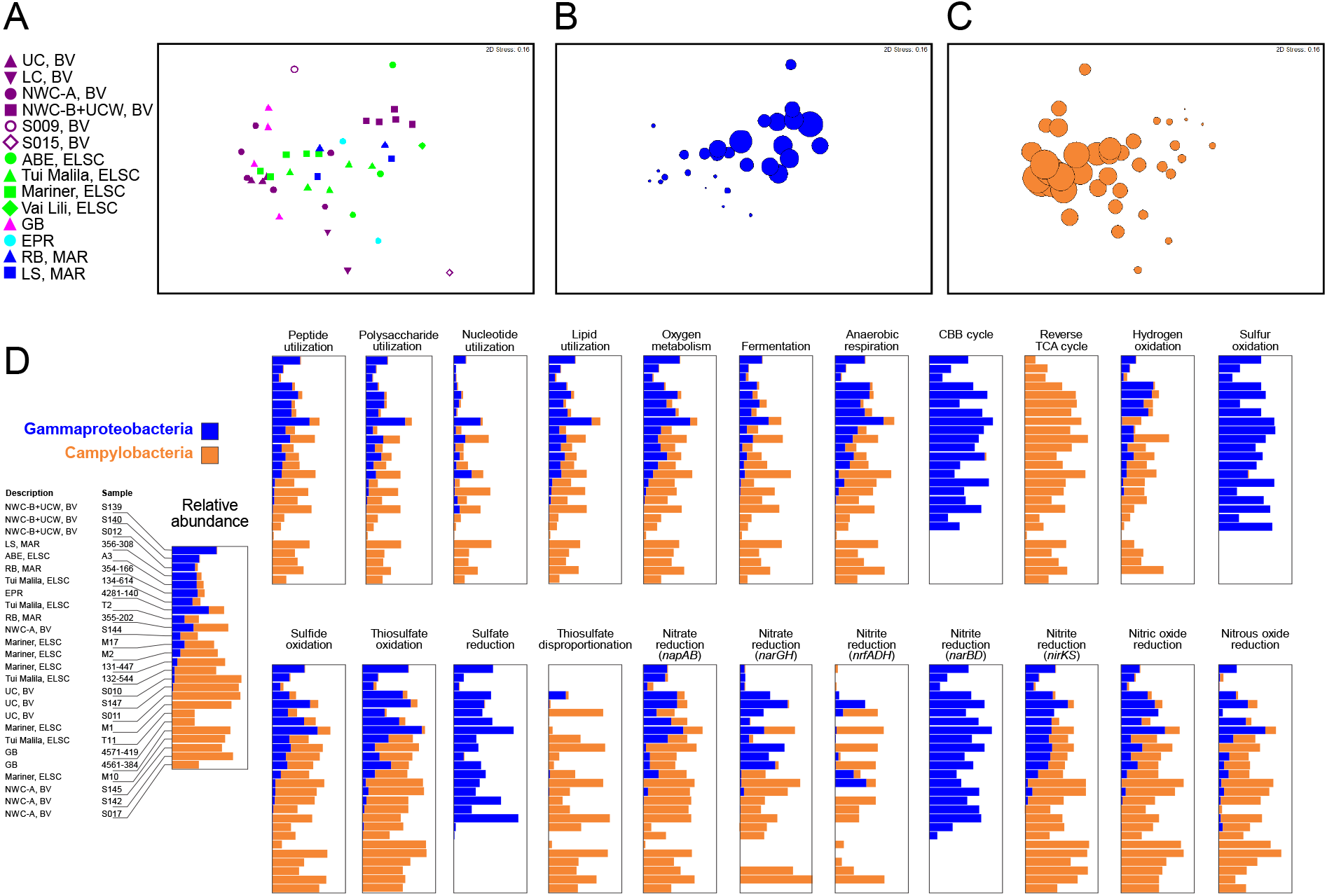
Comparative taxonomic and functional gene abundance of the Campylobacteria and Gammaproteobacteria. NMDS plots were generated using a Bray-Curtis matrix of relative MAG abundance, based on GTDB-assigned taxonomy at the class level. Plots are shown for (A) all sample sites, and for all sample sites with bubbles proportional to the relative abundance of (B) Gammaproteobacteria and (C) Campylobacteria. Comparative functional distribution (D) is also shown for the Gammaproteobacteria and Campylobacteria for the 26 samples that had a summed relative abundance of both Gammaproteobacteria and Campylobacteria of ≥30%. The 22 functions depicted were selected as the Gammaproteobacteria and Campylobacteria accounted for an average of ≥20% of the total abundance for each function across the metagenomes.

The covariation of the Campylobacteria and Gammaproteobacteria in our data also coincided with genes for key functional processes associated with these taxa (Fig. 10). Thus, the overall ecological function contributed by the Campylobacteria and Gammaproteobacteria to the community at all sites was similar, but carried out by either one; viz., same guild different taxa. For example, relative gene abundance for 15 out of 22 broadly distributed functions – including heterotrophy associated with various organic carbon compounds, respiration of oxygen and nitrogen compounds, and oxidation of reduced sulfur compounds – tracked the relative abundance of each group. However, genes for some functions were exclusively represented by either group (Fig. 10, Table S10). For example, marker genes for formaldehyde oxidation, urea utilization and elemental sulfur oxidation were found in the Gammaproteobacteria but were hardly detected in Campylobacteria, while genes associated with thiosulfate disproportionation were attributed almost exclusively to Campylobacteria. In some cases, metabolic analysis also suggested that both Campylobacteria and Gammaproteobacteria had similar metabolic capabilities but encoded different pathways for the same functions. For example, consistent with a previously observed but non-ubiquitous trend [90–93], Campylobacteria mostly encoded genes for the rTCA cycle while the Gammaproteobacteria encoded genes for the CBB cycle. Both taxa also showed the potential for nitrite reduction to ammonia, with more *nrfADH* genes identified in the Campylobacteria and *nirBD* only found in the Gammaproteobacteria.

## Conclusions

From a comparative metagenomic analysis of 38 deep-sea hydrothermal deposits from multiple globally distributed sites, we provide insights into the shared vent-specific lineages and greatly expand the genomic representation of core taxa that have very few, if any, examples in cultivation. Furthermore, we document many novel high quality assembled genomes that were originally only identified from deep-sea vents as 16S rRNA genes. This study sheds light on the metabolic potential and physiological ecology of such taxa. We show that overall, the different communities share similar functions, but differences in the environmental geochemistry between sites select distinct taxonomic guilds. Further, metabolic handoffs in communities provides functional interdependency between populations achieving efficient energy and substrate transformation, while functional redundancy confers higher ecosystem resiliency to perturbations and geochemical fluctuations. In summary, this study provides an integrated view of the genomic diversity and potential functional interactions within high temperature deep-sea hydrothermal deposits and has implications on their biogeochemical significance in mediating energy and substrate transformations in hydrothermal environments.

## Methods

### Sample collection, DNA extraction and sequencing

High-temperature, actively venting deep-sea hydrothermal deposits, a diffuse flow sample, and a water sample were collected from Brothers volcano (2018), the Eastern Lau Spreading Center (2005 and 2015), Guaymas Basin (2009), the Mid-Atlantic Ridge (2008), and the East Pacific Rise (2004 and 2006) as previously described (Flores et al., 2012a, Reysenbach et al., 2020). Expedition details, including identification numbers, research vessels and submersibles utilized for sampling are described in Table S1. Samples were processed [4] and DNA extraction was performed as previously described [4, 8, 25, 96].

### Thermocouple array from Guaymas Basin

The thermocouple array experimental set up from Guaymas Basin in 2009 is described in Teske et al., 2016 [20].

### Metagenomic assembling and binning

Reads from Brothers volcano and ELSC (2015) were quality filtered using FastQC v.0.11.8 (https://www.bioinformatics.babraham.ac.uk/projects/fastqc/) and *de novo* assembled using metaSPAdes v.3.12.0 [97] with the settings “-k 21,33,55,77,99,127 -m 400 --meta”. Reads from ELSC (2005), MAR, EPR and Guaymas Basin were assembled by the Department of Energy, Joint Genome Institute (JGI) using metaSPAdes v.3.11.1 with the settings “-k 33,55,77,99,127 -- only-assembler --meta”. Individual assemblies were generated for each metagenomic dataset. MetaWRAP v.1.2.2 [98] was used to generate metagenome-assembled genomes (MAGs) from each assembly with the settings “--metabat2 --metabat1 --maxbin2”. DAS Tool v.1.0 [99] was then applied to screen the three sets of MAGs generated by MetaWRAP, resulting in consensus MAGs with a minimum scaffold length of 1000 bp.

### Metagenome-assembled genome curation and quality assessment

CheckM v.1.0.7 [100] was used to assess MAG quality and screen for the presence of 16S rRNA genes. Erroneous SSU genes were then removed using RefineM v.0.0.20 [101], which was also used to identify and remove outlier scaffolds with abnormal coverage, tetranucleotide signals, and GC patterns from highly contaminated MAGs. GTDB-Tk v.1.5.0, data release 202 [17] was used to assign taxonomy to each MAG with default settings. SSU sequences from each MAG were then re-parsed and annotated by SINA v.1.2.11 [102]. Scaffolds containing 16S rRNA gene sequences inconsistent with GTDB taxonomic classifications were deemed contaminants and were removed. Selected MAGs were then further refined and manually inspected by VizBin v.1.0.0 [103]. Final MAGs had an estimated ≥50% genome completion and ≤10% contamination, with completeness and contamination rounded to the nearest whole number.

### Iterative Nanoarchaeota MAG curation

As a case study, two MAGs assigned to the Nanoarchaeota (4571-419_metabat1_scaf2bin.008, M10_maxbin2_scaf2bin.065) were iteratively curated, demonstrating that the original MAGs generated by DAS Tool contained large quantities of contaminant contigs that were not recognized by CheckM, given the low abundance of marker genes. Each MAG was visualized using the Anvi’o v.7.1 interactive interface [104], where contigs were divided into subsets based on clustering patterns in Anvi’o. Contigs in each cluster were assigned a putative taxonomy using the Contig Annotation Tool (CAT) [105]. Clusters containing most of the contigs assigned to the Nanoarchaeota were repeatedly sub-sampled and screened using the CAT pipeline until no meaningful correspondence between clustering patterns and assigned taxonomy could be identified (Fig. S5). Contigs in the final clusters were then removed if CAT definitively assigned them to a taxonomic group outside the Nanoarchaeota, while contigs assigned to the Nanoarchaeota and unclassified higher ranks were retained. A third Nanoarchaeota MAG (4281-140_maxbin2_scaf2bin.078) was also identified, but attempted curation using the above workflow revealed the presence of extensive contamination, with only a very small subset of scaffolds confidently assigned to the Nanoarchaeota. CAT analysis of a putative Nanoarchaeota MAG (JGI Bin ID 3300028417_39) separately assembled from the same read set by the JGI as part of the Genomes from Earth’s Microbiomes project [106] also showed very few contigs assigned to the DPANN superphylum and extensive bacterial contamination, suggesting that this particular read set may represent a challenge for commonly utilized binning algorithms. Given the extensive contamination and difficulty identifying a valid Nanoarchaeota MAG of significant size, the 4281-140_maxbin2_scaf2bin.078 was excluded from the MAG dataset submitted to Genbank, so as to avoid contaminating the public database with erroneous information. However, the MAG was included in functional and relative abundance calculations.

### MAG characterization and annotation

Open reading frames (ORFs) were predicted by Prodigal v.2.6.3 [107] with the parameter “-p meta”. ORFs were then annotated by KOfam [108] and custom HMM profiles within METABOLIC v.4.0 [109] and eggNOG-emapper v.2.1.2 [110] with default settings. Transfer RNAs were predicted using tRNAscan-SE 2.0 using the general tRNA model [111]. Genomic properties, including genome coverage, genome and 16S rRNA taxonomy, tRNAs, genome completeness and scaffold parameters were parsed from results that were calculated by CheckM, tRNAscan-SE 2.0 and METABOLIC. Relative genome coverages were normalized by setting each metagenomic dataset size as 100M paired-end reads.

Prior to detailed metabolic analysis, open reading frames from the Gracilibacteria orders BD1-5 and Absconditabacterales, which are known to use genetic code 25 [e.g. 47, 48, 112, 113], were re-called using Prodigal v.2.6.3 as implemented in Prokka v.1.14.6 [114]. An additional MAG from the Gracilibacteria order GCA-2401425 (4559-240_metabat1_scaf2bin.085) was also processed using genetic code 25. Currently, the only other genome in GTDB order GCA-2401425 (Genbank accession NVTB00000000.1) [115] is publicly available in Genbank with ORFs generated using genetic code 11. However, comparative analysis of our GCA-2401425 MAG showed that ORFs called with genetic code 11 were truncated, with an average length of approximately 85 amino acids, while those called with genetic code 25 averaged 277 amino acids in length. ORFs from two additional MAGs from the Paceibacteria (A3_metabat2_scaf2bin.333 and S145_metabat2_scaf2bin.004) were also re-generated in Prokka using genetic code 11. Open reading frames were then annotated in GhostKoala [116].

### Phylogenomic inference

For archaeal phylogenomic tree construction, a concatenated multiple sequence alignment (MSA) was generated in GTDB-Tk using 122 archaeal marker genes (2991 sequences, 5124 columns) [17]. IQ-TREE v.1.6.9 [117] was used to reconstruct the tree with the settings “-m MFP -bb 1000 -redo -mset WAG,LG,JTT,Dayhoff -mrate E,I,G,I+G -mfreq FU -wbtl” (Data S1). The bacterial phylogenomic tree was constructed in a similar manner, using a concatenated MSA of 120 bacterial GTDB marker genes [17]. For each GTDB bacterial phylum, no more than 15 reference genomes from the GTDB r202 database were used (4248 sequences, 5037 columns; Data S2). Additionally, a second bacterial phylogenomic tree was inferred from the same MSA using FastTree v.2.1.8 (WAG, +gamma, SH support; Data S3) [118]. Additional MSAs solely using MAGs from this study were generated for the Archaea (122 marker genes) and Bacteria (120 marker genes) using the GTDB-Tk identify and align commands [17]. FastTree v.2.1.10 (parameter: --gamma) was used to infer the phylogenomic trees, as implemented in GTDB-Tk (Data S4, S5; formatted trees available online at https://itol.embl.de/shared/alrlab).

A tree was constructed in GTDB-Tk (parameter: --gamma) using MAGs assigned to the Patescibacteria, along with recently described *Cand*. Vampirococcus lugosii [47] and *Cand*. Absconditicoccus praedator [48], and the GTDB r202 bacterial tree-building dataset. A phylogenomic tree of the Chloroflexota was also generated by extracting a concatenated MSA of Chlorofexota MAGs from the entire bacterial MSA. IQ-TREE v.2.1.4 [119] was used to reconstruct the tree with the settings “-m TESTMERGE -bb 1000 -bnni”. An outgroup genome (GCA_007123655.1) was added to reroot the phylogenomic tree. Final trees were visualized using Interactive Tree of Life (iTOL) v.6 [120].

### Taxonomic assignment

Initial taxonomy was assigned to each MAG using the GTDB-Tk classify pipeline. In rare instances where there were discrepancies between the class-level (Archaea) or phylum-level taxonomy (Bacteria) assigned by GTDB-Tk and phylogenetic tree topology, we deferred to tree topology. In the Bacteria, topological taxonomic assignments were only used if confirmed by both trees. MAGs that were not assigned to a known genus by GTDB-Tk were compared to their closest relatives in this study using average amino acid identity (AAI) matrices generated in CompareM v.0.1.2 (https://github.com/dparks1134/CompareM). MAGs were assigned to novel genera using cutoffs provided by Konstantinidis et al. [18], and MAGs assigned the taxonomic status “unclassified” were automatically assigned to a novel genus.

### Trophic and energy metabolism analysis

Functional genes were first characterized by METABOLIC [109]. Additional peptide utilization genes were characterized using the MEROPS database release 12.3 [121], and additional polysaccharide utilization genes were identified using dbCAN2 (2020-04-08) and the CAZy (2021-05-31) database [122, 123]. Cellular localization of peptidases/inhibitors, gene calls identified by the CAZy database, and predicted extracellular nucleases were verified using PSORTb v.3.0 [124]. Functional annotations for protein, polysaccharide, nucleic acid and lipid utilization were derived in part from previous publications [125, 126]. Iron cycling genes and hydrogenase genes were characterized based on HMMs directly obtained or indirectly parsed from FeGenie [127] and HydDB [75].

For each of these trophic and energy metabolisms, the number of functional gene calls in each genome were calculated using two different scenarios: 1) the presence of any marker gene in the complex/pathway was treated as the presence of the whole function (indicated as C), and the highest number of gene calls for an individual gene in the complex was taken to be the number of pathway ‘hits’ in the MAG. 2) Stand-alone genes that were not part of a large complex or functional pathway (indicated as A) were treated as individual accumulative gene calls for their particular function. In specific cases, marker genes were manually verified using phylogenetic trees and by inspecting operon arrangements (see below). To calculate functional abundance, all genomes were included in the analysis. Functional abundance was then calculated by multiplying normalized genome coverage (100M reads/sample) by the number of functional gene calls for each sample. For visualization, functional abundance was then log-transformed and used to generate heatmaps with the R package pheatmap v.1.0.12 (settings: clustering_method = ward.D2). Combined functional heatmaps were also generated by summing values within larger functional groups.

To avoid potential mis-annotation by the automated methods described above, phylogenetic trees were constructed to validate predicted protein sequences for dissimilatory sulfite reductase (Dsr; Fig. S9), methyl-coenzyme M reductase subunit alpha (McrA; Fig. S10) and sulfur dioxygenase (Sdo; Fig. S11). Based on current understanding, two metabolic directions are possible for the Dsr protein: reductive Dsr, which catalyzes the reduction of sulfite to sulfide, and oxidative (or reverse) Dsr, which converts elemental sulfur oxidation to sulfite [128]. Paired DsrAB proteins were first identified in all MAGs using in-house Perl scripts. In cases where Dsr subunits were duplicated, one set of paired DsrAB proteins was manually selected. A concatenated protein alignment was then generated for DsrAB proteins from the MAGs and reference sequences using MAFFT v.7.310 [129], and the alignment was trimmed using trimAl v.1.4.rev15 [130] with the parameter “-gt 0.25”. A phylogenetic tree was then constructed in IQ-TREE with settings “-m MFP -bb 1000 -redo -mset WAG,LG,JTT,Dayhoff -mrate E,I,G,I+G - mfreq FU -wbtl” (Fig. S9). Reductive and oxidative DsrAB proteins were identified based on placement in the phylogenetic tree.

Predicted proteins for McrA were first identified using the TIGR03256 HMM. Presumed false gene calls were then manually removed, including those identified in bacterial MAGs and non-methanogenic/anaerobic methanotrophic archaeal MAGs with high sequence coverage. An alignment was constructed in MAFFT v.7.310 [129] using the remaining McrA protein sequences, together with reference genes recovered from methanogens, anaerobic methanotrophs, and short-chain alkane oxidizing Archaea from the Bathyarchaeia, Helarchaeales, *Syntrophoarchaeum* and *Polytropus* [11, 12, 131, 132]. Alignment trimming and phylogenetic tree inference were performed as described above.

Sulfur dioxygenase (Sdo) proteins were predicted using the “sulfur_dioxygenase_sdo” HMM [109]. Alignment, trimming and construction of the phylogeny were performed as described above. Positive Sdo calls were identified using two conserved amino acid residues (Asp196 and Asn244 of hETHE1, NCBI accession NP_055112) that are specific to Sdo in comparison with other metallo-β-lactamase superfamily members [133].

### Statistical analysis

The relative abundance of MAGs in this study was calculated for each sample using normalized read coverage (set to 100M reads) expressed as a percentage. Bray-Curtis similarity matrices were then generated from relative abundance data at various taxonomic ranks, and nonmetric multidimensional scaling (NMDS) plots were generated from the matrices using PRIMER v.6.1.13 [134].

### Data availability

Metagenome reads are publicly available in the Sequence Read Archive (Table S1), and MAGs generated in this study are available in NCBI Genbank (BioProject PRJNA821212, Table S2).

## Supporting information

Additional File 8: Supplementary Discussion

## Acknowledgements

We thank the crew of the R/V *Roger Revelle*, R/V *Atlantis*, R/V *Thomas G. Thompson*, HOV *Alvin*, and the ROV *Jason* for assistance in collecting the samples. Many thanks to the many students who over the years helped extract the DNA, and to MK Tivey for thoughtful comments to the manuscript. This work was funded by the US-National Science Foundation grants OCE-0728391, OCE-0937404, OCE-1558795 to A.-L.R, and OCE-2049478 and DBI-2047598 to K.A. We thank the Department of Energy Joint Genome Institute (Community Science Program award 339, lead Peter Girguis) for sequencing several of the samples.

## Additional Supplementary Files

### Additional File 1. Supplementary Figures (.pdf)

**Fig. S1**. Geographic distribution of deep-sea hydrothermal vent sampling locations. The number of samples collected in each region is shown with *n* values. **Fig. S2**. Deep-sea hydrothermal vent photographs from ELSC, EPR, MAR and Guaymas Basin. **Fig. S3**. Comparison between the number of medium- to high-quality MAGs recovered in each metagenomic assembly and the number of reads that passed quality control measures. Metagenomic assemblies are ordered by (A) increasing MAG count and (B) increasing read count. **Fig. S4**. NMDS plot showing the taxonomic diversity of Brothers volcano MAGs, based on normalized relative abundance. Clustering patterns show a high degree of similarity to NMDS plot clustering previously reported in Reysenbach et al., 2020. **Fig. S5**. Anvi’o plot showing the cluster of scaffolds (blue) predominantly corresponding to the Nanoarchaeota in M10_maxbin2_scaf2bin.065. Analysis with CAT revealed three additional contaminating scaffolds which were removed, bringing the final scaffold count to 149, with an estimated 47% completion by CheckM. Scaffold clusters that were removed (pink; 972 scaffolds) were largely assigned to taxonomic groups outside the Nanoarchaeota by CAT and had a low number of marker genes, as estimated by CheckM (6.99% completion, 0.29% contamination). **Fig. S6**. Predicted cell metabolism diagrams for the putative new phyla (A) JALSQH01 (3 MAGs) and (B) JALWCF01 (13 MAGs). Functions (F) and modules (M) were identified using METABOLIC (Table S5). Solid lines indicate the presence of a module or function, while dashed lines and a “p” in parentheses indicate that a module or function was only present sporadically (<50% of MAGs). Modules and functions not identified in any MAGs are shown with dashed lines and gray labels. **Fig. S7**. Normalized relative abundance of GTDB classes, expressed as a percentage. Classes depicted comprise ≥16% of the relative MAG abundance in at least one assembly. **Fig. S8**. Maximum-likelihood GTDB-Tk concatenated protein tree showing members of the Patescibacteria, used to generate Fig. 5A. Lineages outside the Patescibacteria are shown as a collapsed triangle, and MAGs from this study are indicated in bold type. Filled circles represent SH-like branch support (0.8-1.0), and the scale bar shows 0.5 substitutions per amino acid. **Fig. S9**. Concatenated dissimilatory sulfite reductase (DsrAB) protein phylogenetic tree. Only the nodes with ultrafast bootstrap (UFBoot) support values over 90% were labeled with black dots. This tree included both reductive DsrAB (for reductive dissimilatory sulfite reduction to sulfide) and oxidative DsrAB (for dissimilatory sulfur oxidation to sulfite). For collapsed clades in the oxidative DsrAB clade (labeled in blue), the DsrAB call numbers and DsrAB-containing MAG numbers were labeled in square brackets. The total number for both reductive DsrAB calls and reductive DsrAB-containing MAG numbers and oxidative DsrAB calls and oxidative DsrAB-containing MAG numbers were labeled accordingly on the side of the tree. Note that one genome can have multiple paired DsrAB calls. **Fig. S10**. Phylogenetic protein tree of methyl coenzyme M reductase subunit alpha (McrA). Ultrafast bootstrap support values (>90%) are shown with filled circles. Clades comprised of predicted butane oxidation (Butane clade), X-alkane oxidation (X-alkane clade) and anaerobic methanotrophy-associated (ANME-1 and -2) McrA amino acid sequences are highlighted, and the three predicted McrA sequences from the Archaeoglobi are shown in red. **Fig. S11**. Sdo (sulfur dioxygenase) phylogenetic protein tree. Only the nodes with ultrafast bootstrap (UFBoot) support values over 90% were labeled with black dots. The positive Sdo sequences that were checked by two conservative amino acid residues were labeled yellow in the tree. Three positive Sdo clades (including ETHE1, Sdo, and Blh) were labeled yellow; the numbers of positive Sdo sequences, non-Sdo sequences, and Sdo reference sequences were labeled accordingly. Other unannotated clades and non-Sdo clades (including metallo-beta-lactamase, GloB1, and GloB2) all contained non-Sdo sequences. **Fig. S12**. Relative abundance of GTDB-assigned MAG taxa at Guaymas Basin. Abundances are shown (A) for all taxa at the genus level, and (B) for the Archaea at the order level, using read coverage normalized to 100M reads per sample and expressed as a percentage of MAG reads per sample. Relative abundances were averaged for the two samples from the six-day thermocouple array (4561-380 and 4561-384).

### Additional File 2. Supplementary Tables (.xlsx)

**Table S1**. Sample metadata including location, year, research vessel, number of metagenome reads and accession numbers. **Table S2**. MAG genome properties, accession numbers and taxonomic classifications. Taxonomy was assigned using GTDB-Tk, and mis-classified MAGs were taxonomically re-assigned at the phylum level (Bacteria) and class level (Archaea) using curated archaeal and bacterial phylogenetic trees. Genome quality statistics are based solely on completion (high-quality draft, >90%; medium-quality draft, >50%). **Table S3**. Average amino acid identity (AAI) matrices for the (A) Bacteria and (B) Archaea. Matrices are grouped by GTDB taxonomy and include MAGs that could not be assigned to a known genus by GTDB-Tk. **Table S4**. Relative abundance of GTDB taxa by site, based on read coverage of MAGs normalized to 100M reads per sample. MAG coverage for each site was summed and expressed as a percent. **Table S5**. METABOLIC-G results for JALSQH01 (3 MAGs) and JALWCF01 (13 MAGs). In the summary rows for JALSQH01 and JALWCF01, functions and modules are listed as “present” if identified in ≥50% of all MAGs, “partially present” if found in <50% of the MAGs, and “absent” if undetected in the MAGs. **Table S6**. Selected functional genes found in Patescibacteria MAGs, based on annotation with GhostKOALA. KEGG module numbers are shown in parentheses. **Table S7**. Functional genes identified in selected high-completeness (≥80%) MAGs from the Chloroflexota. (A) Genes are marked as present (1; green highlight) or not detected (0) in individual MAGs. (B) The proportion of high-completeness MAGs in six GTDB orders that encode functional genes is also shown, with proportions ≥50% highlighted in green. **Table S8**. Identification and distribution of functional genes in this study. (A) The HMMs, MEROPS peptidases, and CAZymes used to identify functional genes. Gene call numbers were calculated using the component (C) or accumulative (A) methods described in the methods. Genes requiring manual validation (M) are indicated. (B) Functional gene abundance, calculated as described in the methods. **Table S9**. Percentage of MAGs in phylogenetic clusters that encode core metabolic genes. Unless otherwise indicated, Archaea are shown at the class level, and Bacteria are shown at the phylum level. Genes were detected using METABOLIC, with additional validation steps for oxidative and reductive Dsr, Sdo, PmoA and McrA. **Table S10**. Comparative (A) relative abundance and (B) functional gene abundance for the Gammaproteobacteria and Campylobacteria, used to generate **Fig. 10**.

### Additional File 3. Data S1 (.nwk)

Newick format archaeal concatenated protein phylogenetic tree, including both MAGs and GTDB reference genomes.

### Additional File 4. Data S2 (.nwk)

Newick file of bacterial concatenated protein phylogenetic tree including MAGs and GTDB reference genomes, generated using IQ-TREE.

### Additional File 5. Data S3 (.nwk)

Concatenated protein phylogenetic tree of bacterial MAGs and GTDB reference genomes, generated with FastTree (Newick format).

### Additional File 6. Data S4 (.nwk)

MAG-only bacterial concatenated phylogenetic protein tree in Newick format.

### Additional File 7. Data S5 (.nwk)

Concatenated protein phylogeny of archaeal MAGs in Newick format.

### Additional File 8. Supplementary Discussion (.docx)

