## Additional File 1: Supplementary Figures for "Global patterns of diversity and metabolism of microbial communities in deep-sea hydrothermal vent deposits"

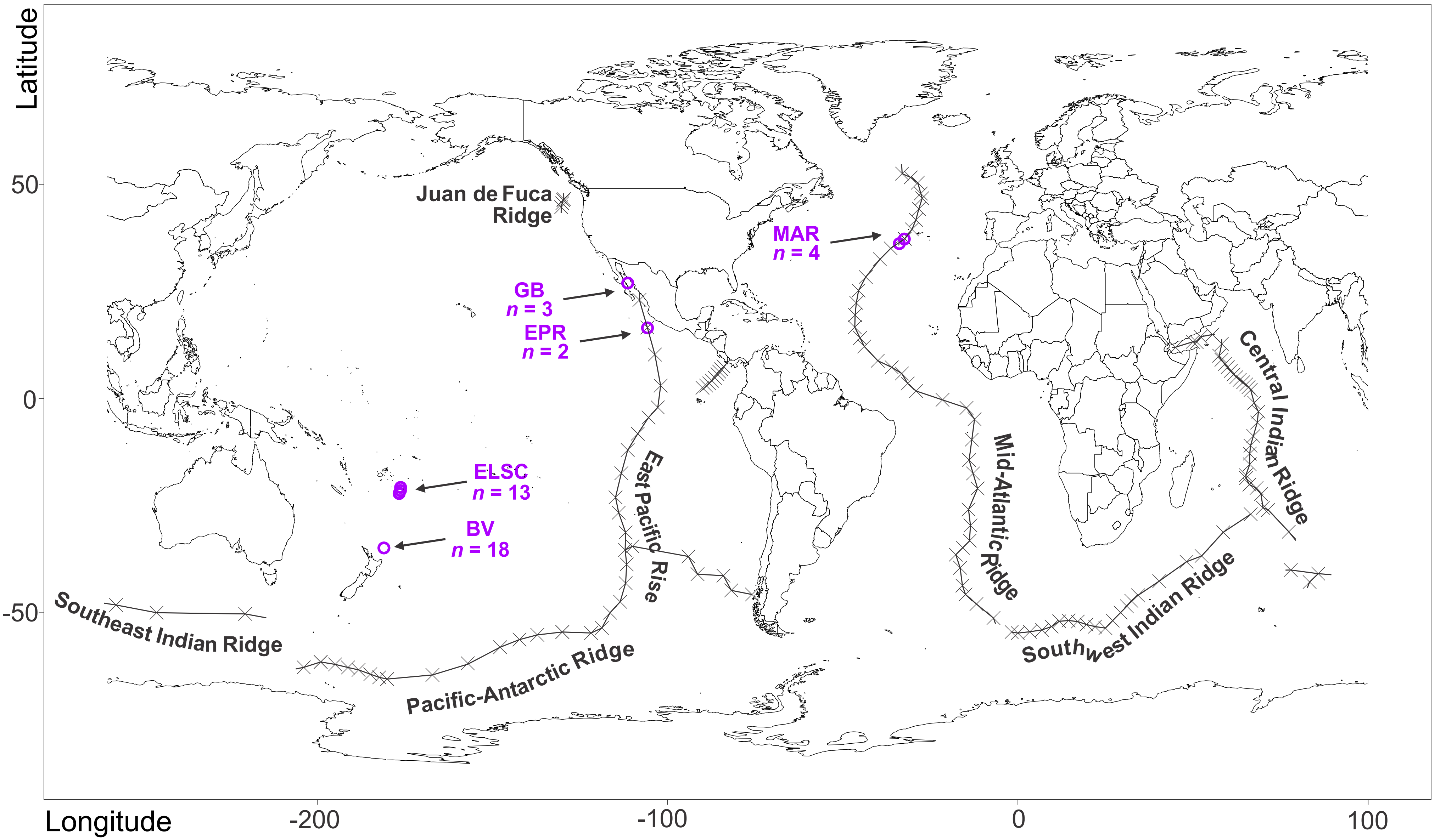


**Fig. S1.** Geographic distribution of deep-sea hydrothermal vent sampling locations. The number of samples collected in each region is shown with *n* values.

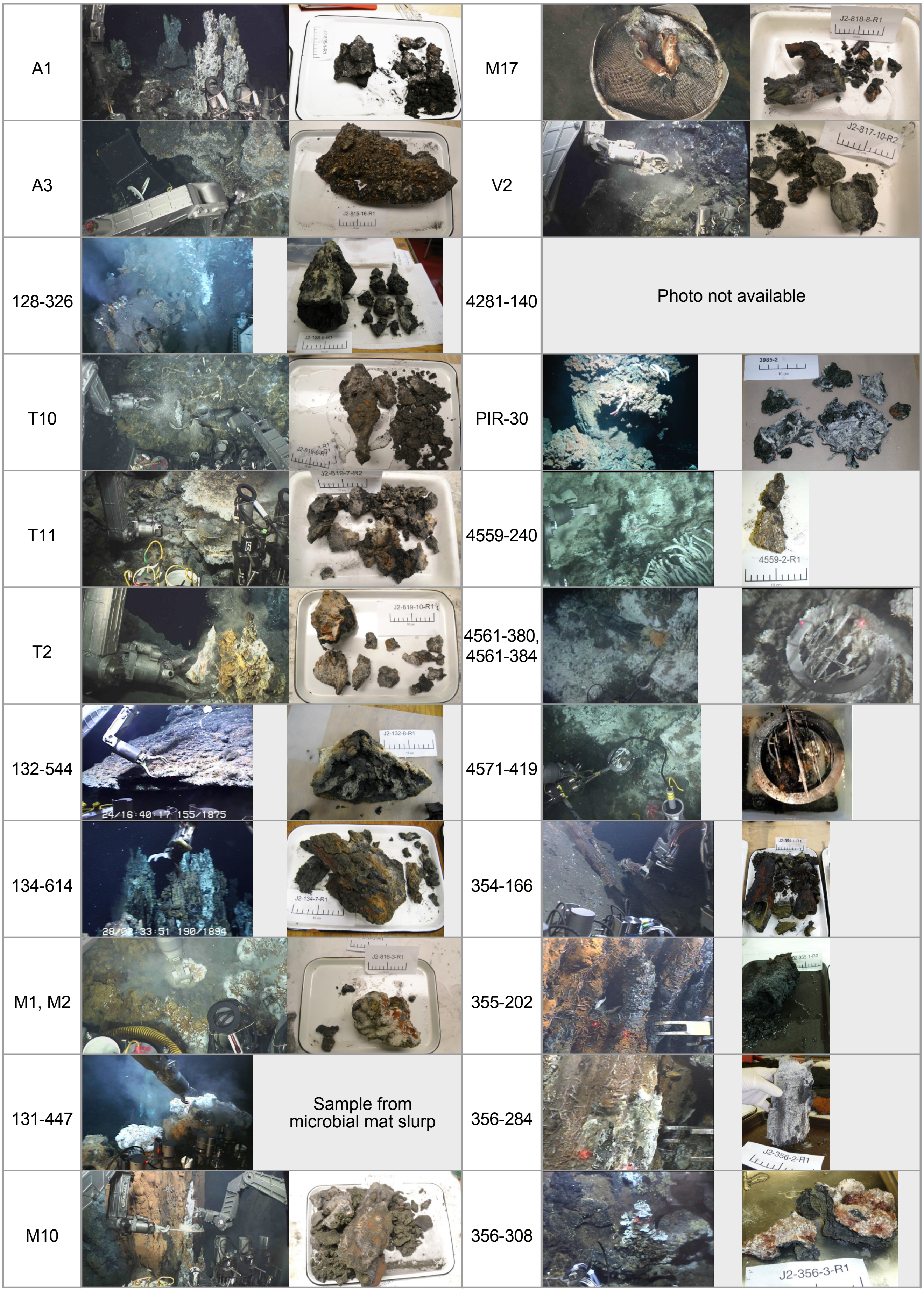


**Fig. S2.** Deep-sea hydrothermal vent photographs from ELSC, EPR, MAR and Guaymas Basin.

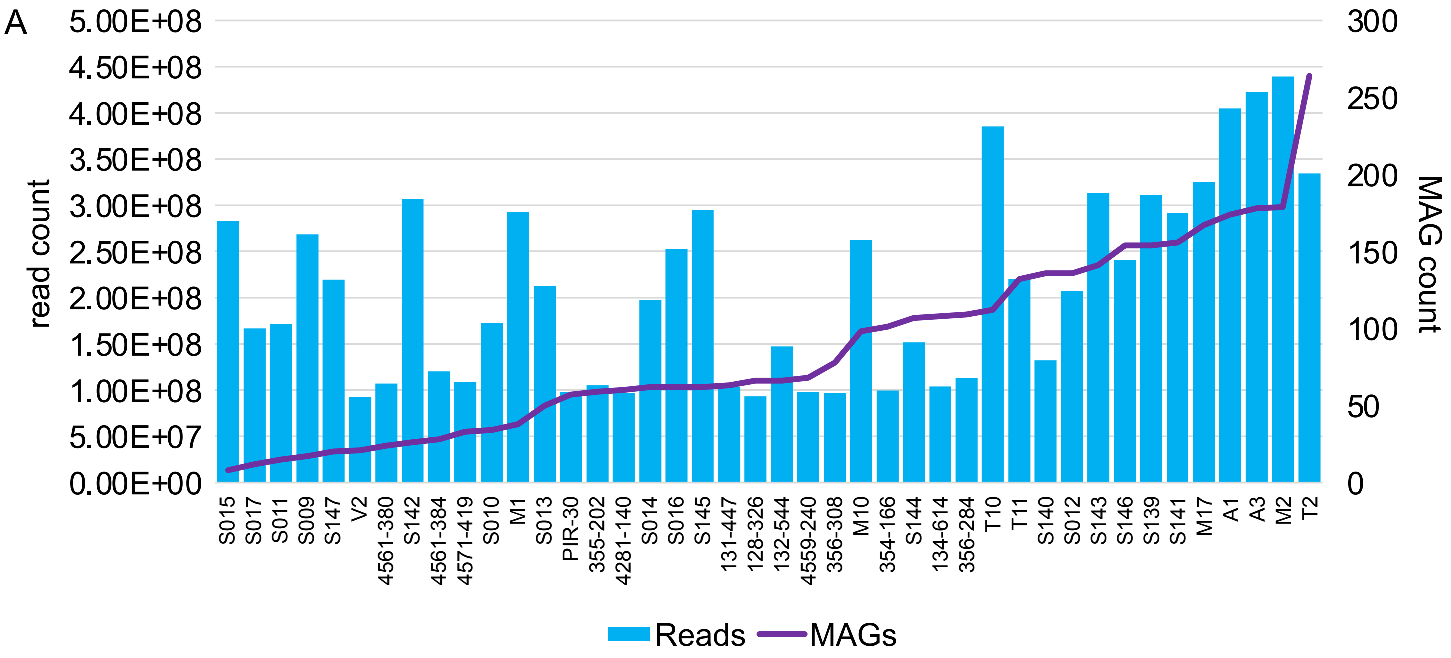


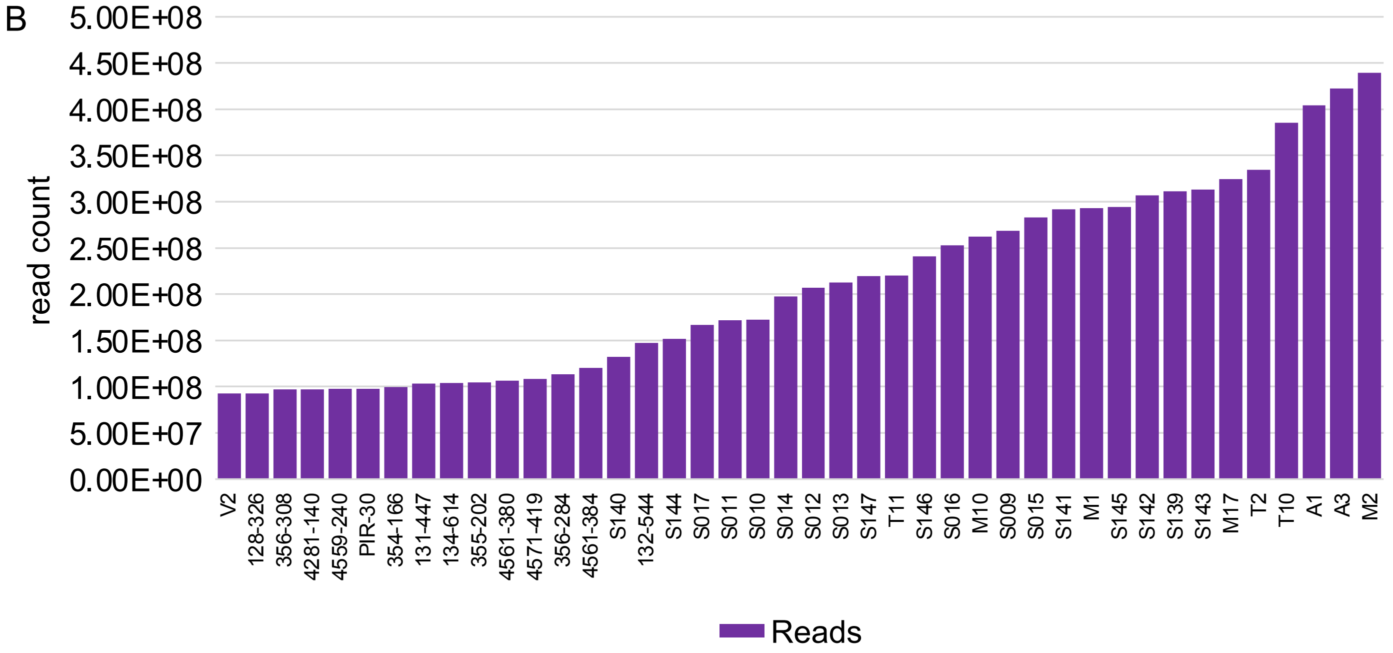


**Fig. S3.** Comparison between the number of medium- to high-quality MAGs recovered in each metagenomic assembly and the number of reads that passed quality control measures. Metagenomic assemblies are ordered by (A) increasing MAG count and (B) increasing read count.

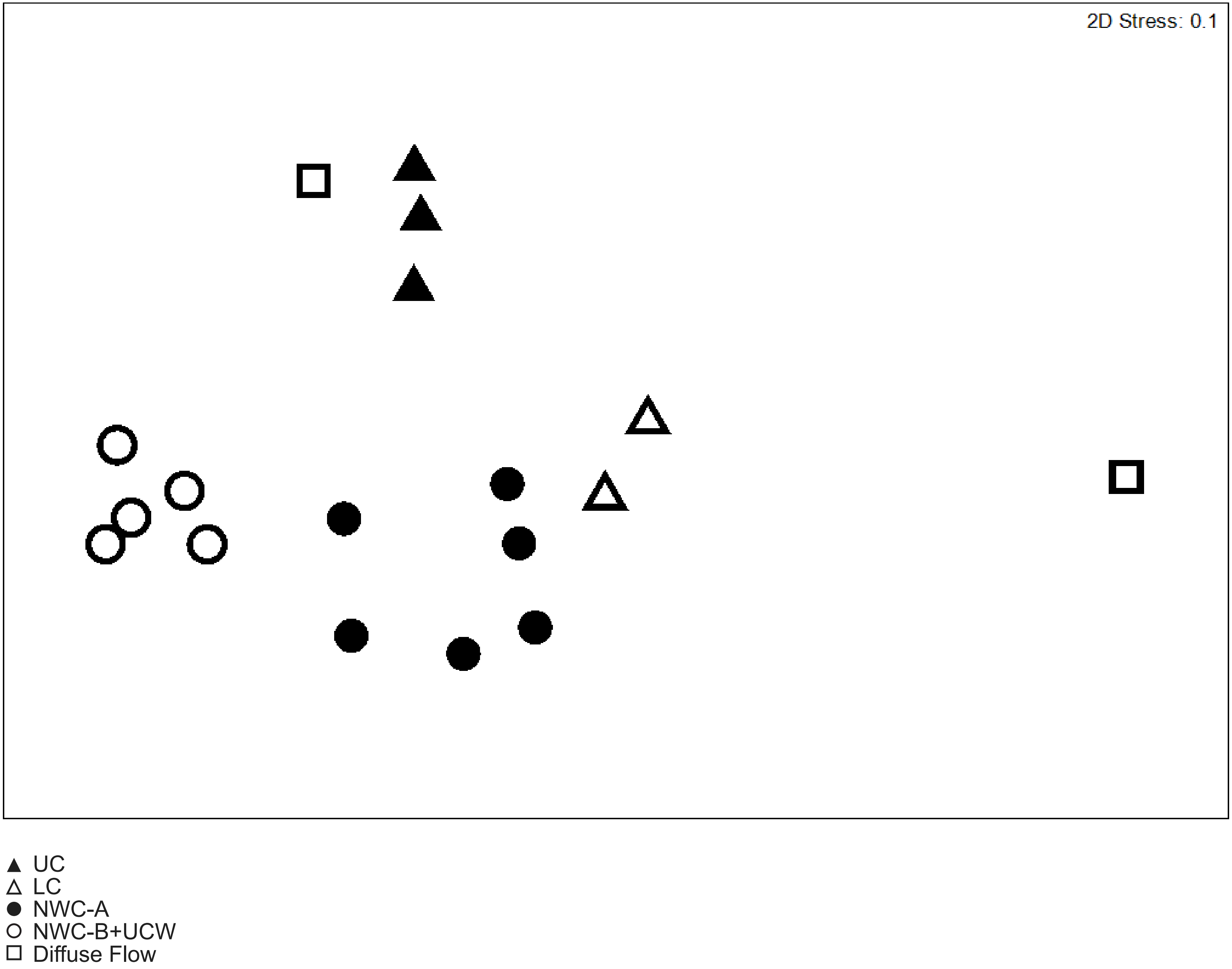


**Fig. S4.** NMDS plot showing the taxonomic diversity of Brothers volcano MAGs, based on normalized relative abundance. Clustering patterns show a high degree of similarity to NMDS plot clustering previously reported in Reysenbach et al., 2020.

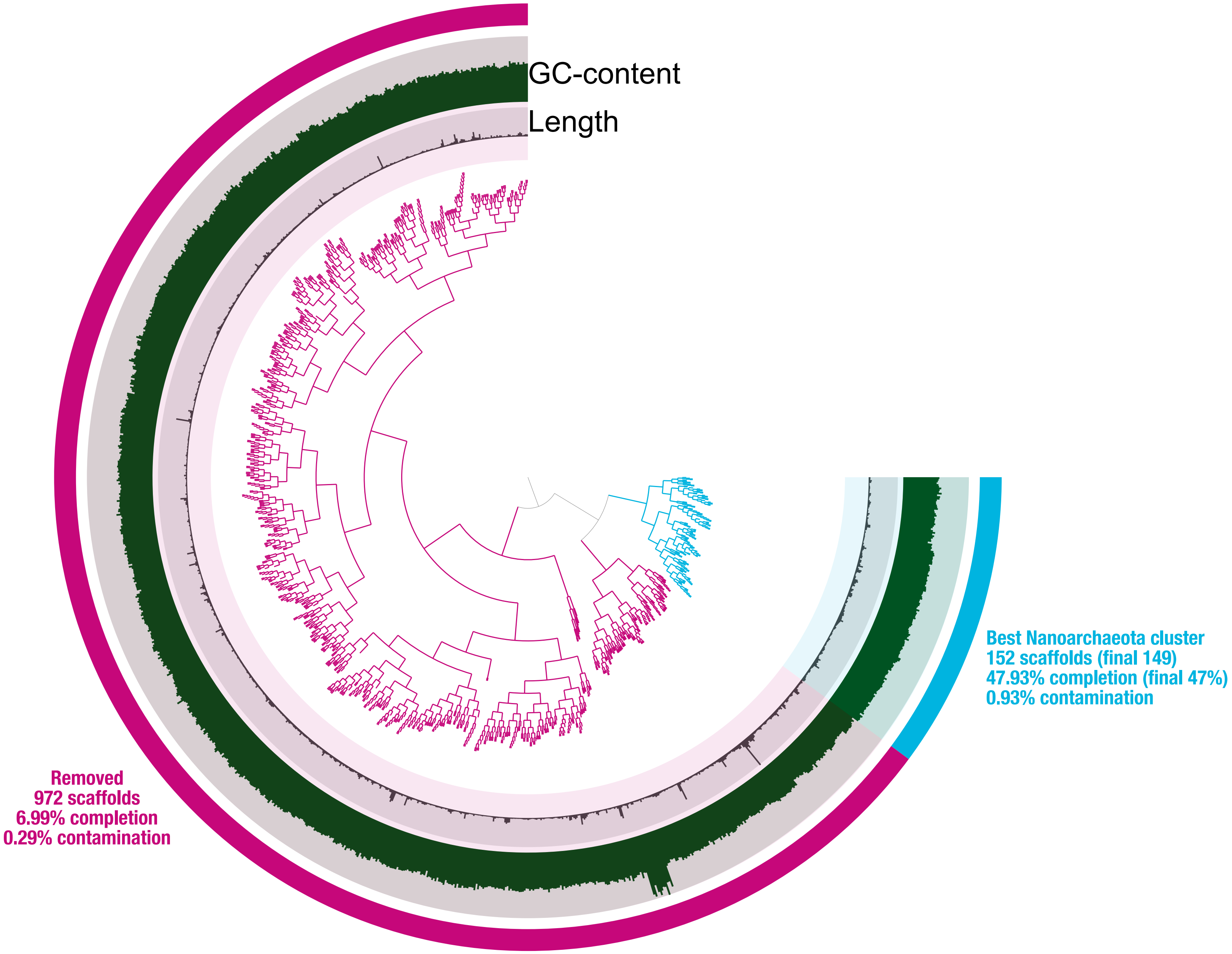


**Fig. S5.** Anvi'o plot showing the cluster of scaffolds (blue) predominantly corresponding to the Nanoarchaeota in M10_maxbin2_scaf2bin.065. Analysis with CAT revealed three additional contaminating scaffolds which were removed, bringing the final scaffold count to 149, with an estimated 47% completion by CheckM. Scaffold clusters that were removed (pink; 972 scaffolds) were largely assigned to taxonomic groups outside the Nanoarchaeota by CAT and had a low number of marker genes, as estimated by CheckM (6.99% completion, 0.29% contamination).

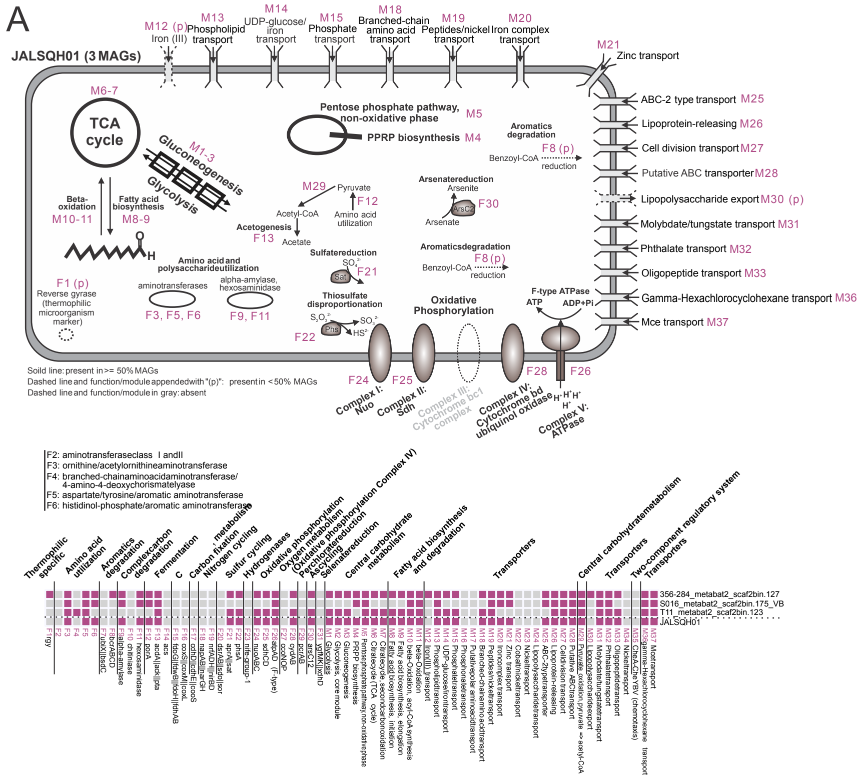


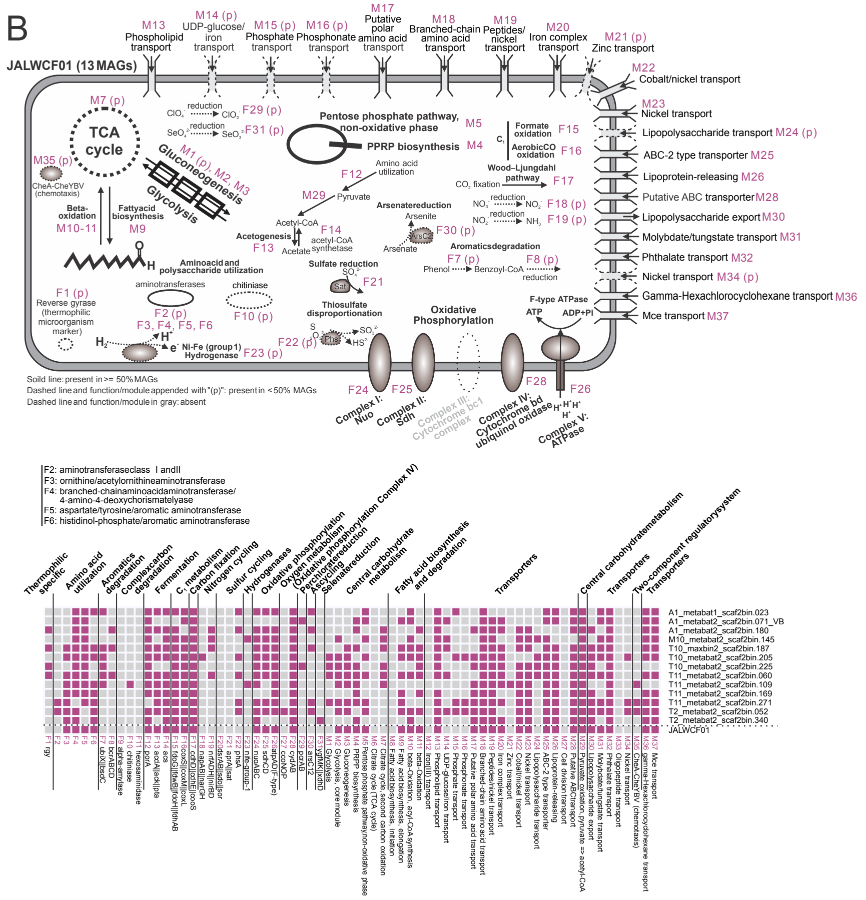


**Fig. S6.** Predicted cell metabolism diagrams for the putative new phyla (A) JALSQH01 (3 MAGs) and (B) JALWCF01 (13 MAGs). Functions (F) and modules (M) were identified using METABOLIC (Table S5). Solid lines indicate the presence of a module or function, while dashed lines and a “p” in parentheses indicate that a module or function was only present sporadically (<50% of MAGs). Modules and functions not identified in any MAGs are shown with dashed lines and gray labels.

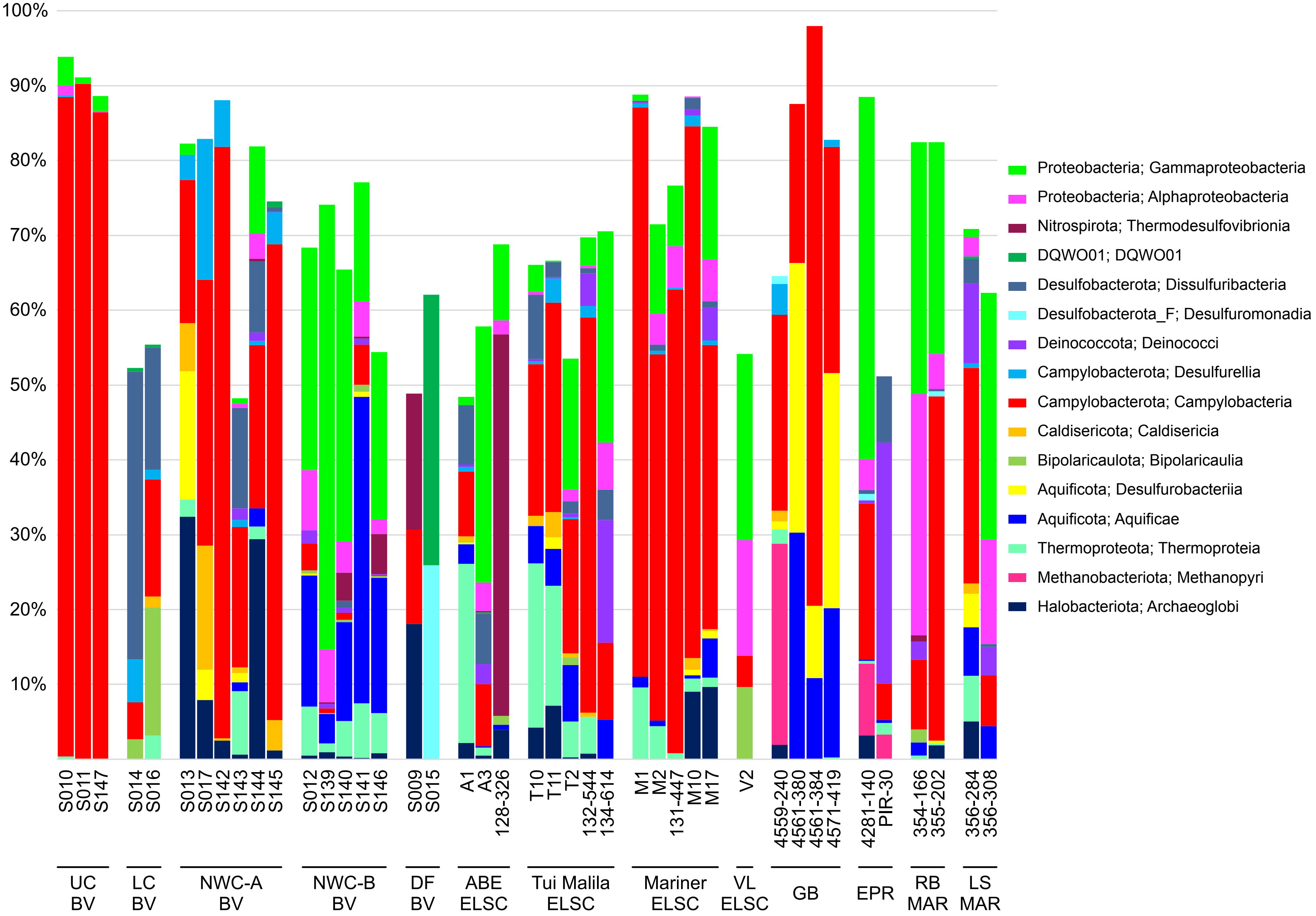


**Fig. S7.** Normalized relative abundance of GTDB classes, expressed as a percentage. Classes depicted comprise ≥16% of the relative MAG abundance in at least one assembly.

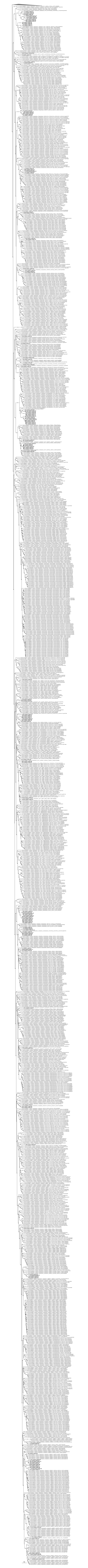

**Fig. S8.** Maximum-likelihood GTDB-Tk concatenated protein tree showing members of the Patescibacteria, used to generate Fig. 5A. Lineages outside the Patescibacteria are shown as a collapsed triangle, and MAGs from this study are indicated in bold type. Filled circles represent SH-like branch support (0.8-1.0), and the scale bar shows 0.5 substitutions per amino acid.

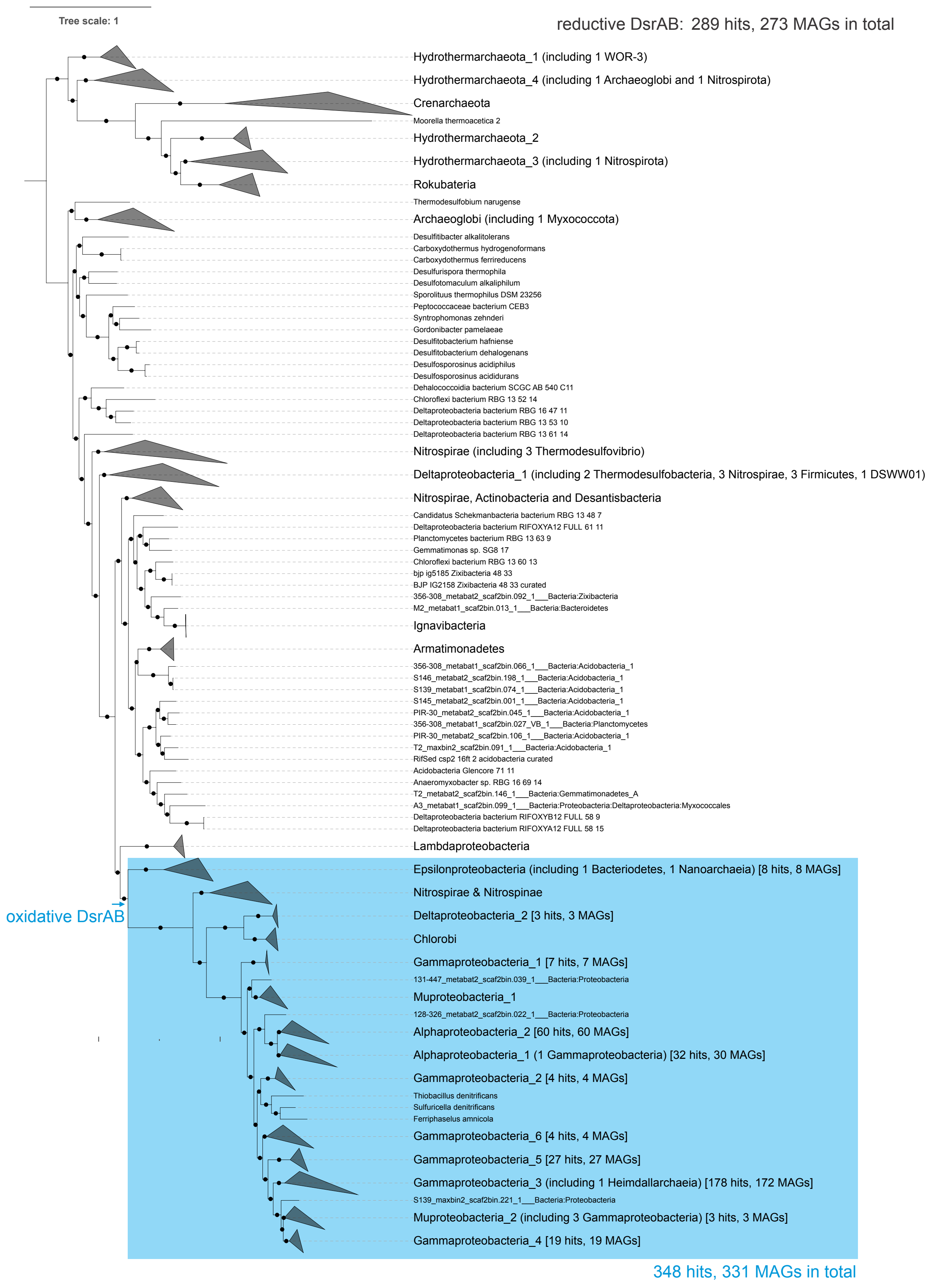


**Fig. S9.** Concatenated dissimilatory sulfite reductase (DsrAB) protein phylogenetic tree. Only the nodes with ultrafast bootstrap (UFBoot) support values over 90% were labeled with black dots. This tree included both reductive DsrAB (for reductive dissimilatory sulfite reduction to sulfide) and oxidative DsrAB (for dissimilatory sulfur oxidation to sulfite). For collapsed clades in the oxidative DsrAB clade (labeled in blue), the DsrAB call numbers and DsrAB-containing MAG numbers were labeled in square brackets. The total number for both reductive DsrAB calls and reductive DsrAB-containing MAG numbers and oxidative DsrAB calls and oxidative DsrAB-containing MAG numbers were labeled accordingly on the side of the tree. Note that one genome can have multiple paired DsrAB calls.

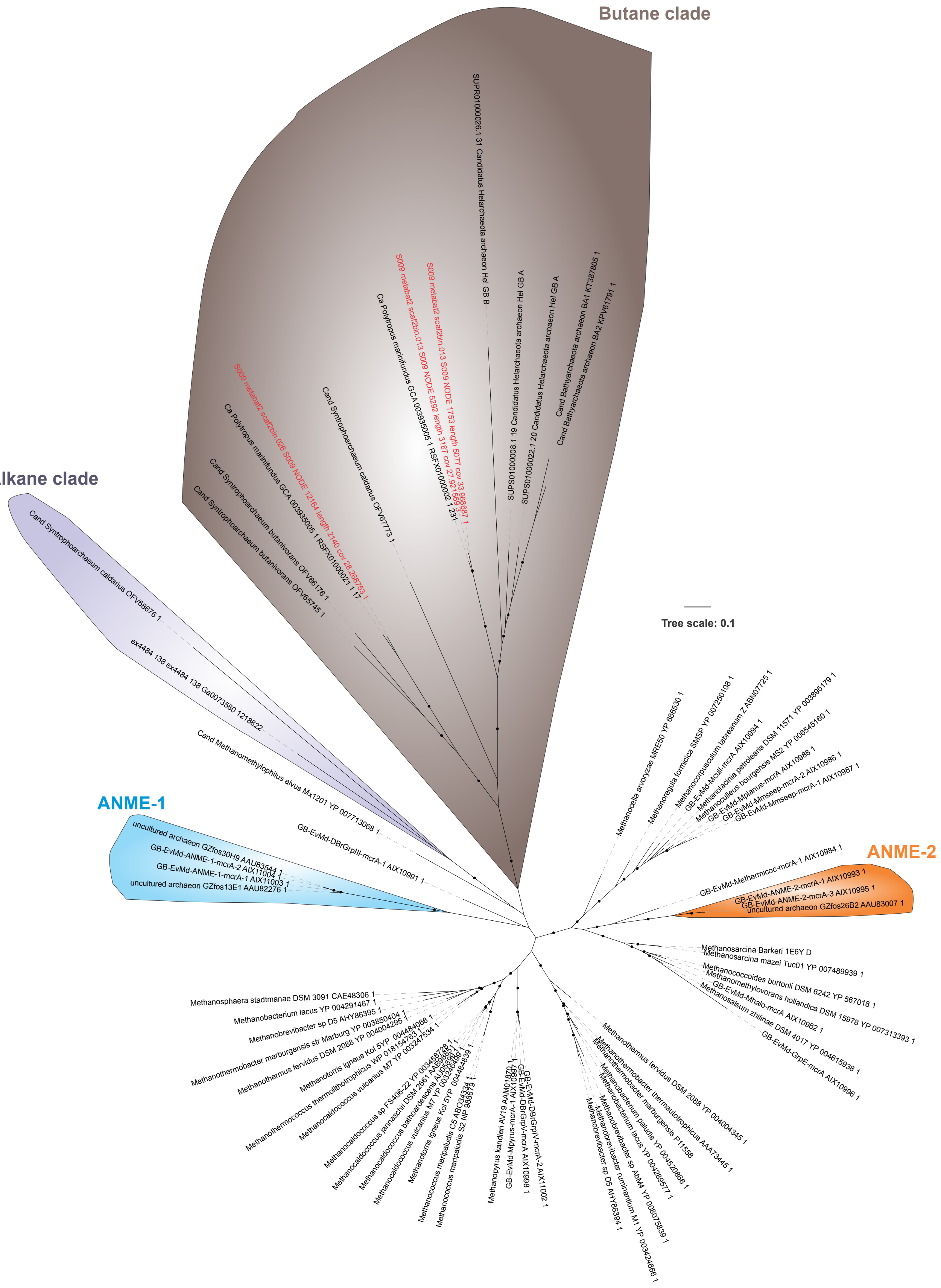


**Fig. S10.** Phylogenetic protein tree of methyl coenzyme M reductase subunit alpha (McrA). Ultrafast bootstrap support values (>90%) are shown with filled circles. Clades comprised of predicted butane oxidation (Butane clade), X-alkane oxidation (X-alkane clade) and anaerobic methanotrophy-associated (ANME-1 and -2) McrA amino acid sequences are highlighted, and the three predicted McrA sequences from the Archaeoglobi are shown in red.

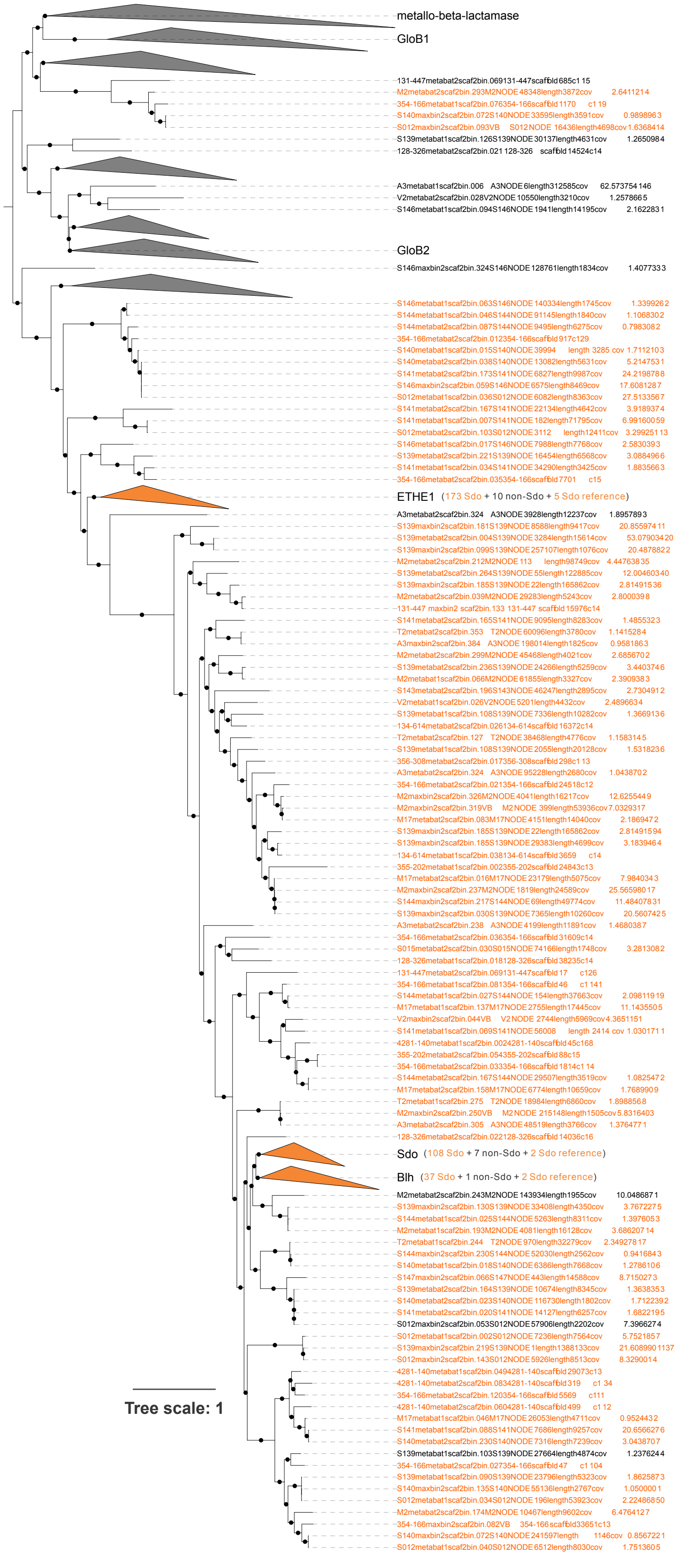


**Fig. S11.** Sdo (sulfur dioxygenase) phylogenetic protein tree. Only the nodes with ultrafast bootstrap (UFBoot) support values over 90% were labeled with black dots. The positive Sdo sequences that were checked by two conservative amino acid residues were labeled yellow in the tree. Three positive Sdo clades (including ETHE1, Sdo, and Blh) were labeled yellow; the numbers of positive Sdo sequences, non-Sdo sequences, and Sdo reference sequences were labeled accordingly. Other unannotated clades and non-Sdo clades (including metallo-beta-lactamase, GloB1, and GloB2) all contained non-Sdo sequences.

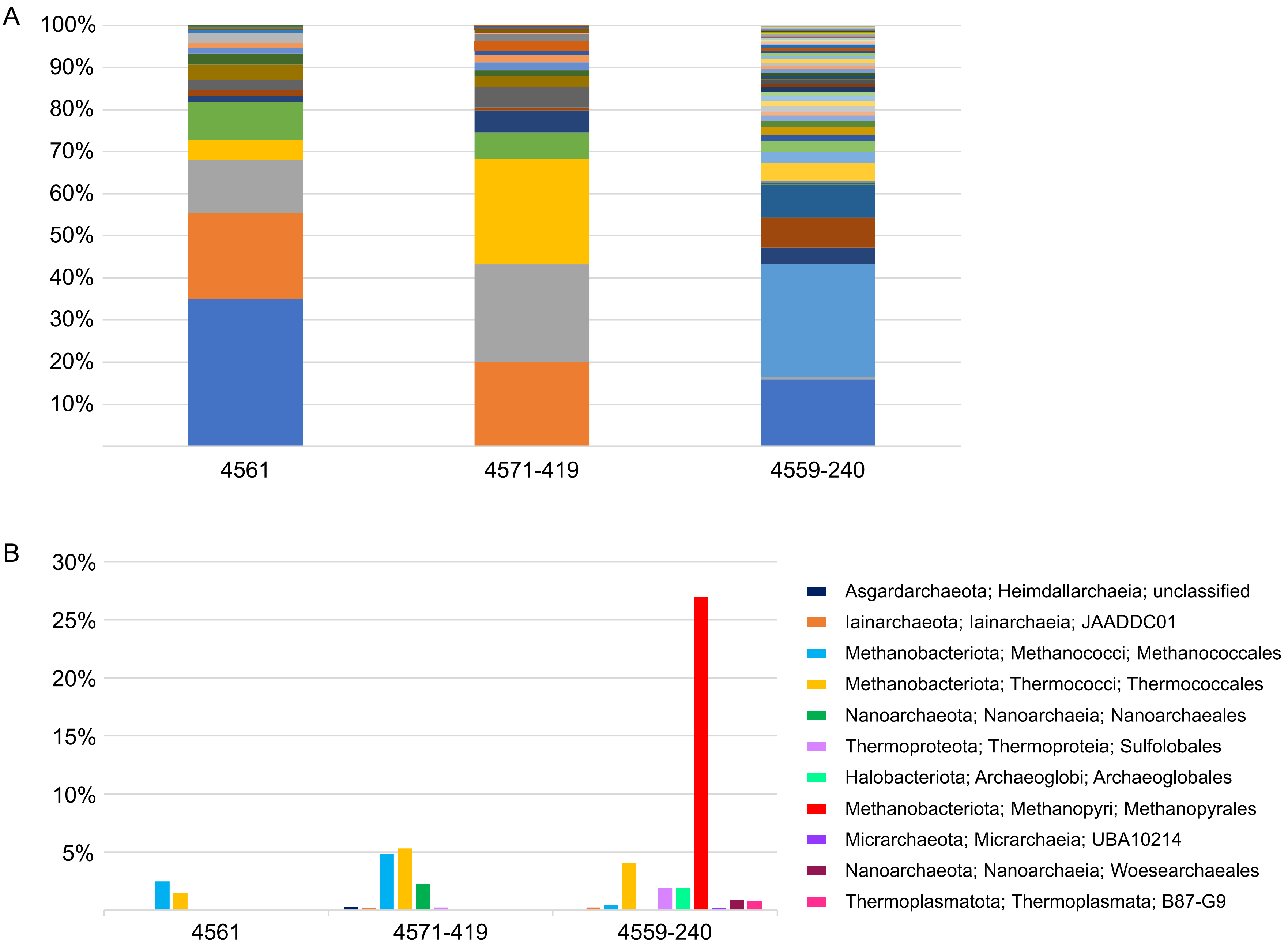


**Fig. S12.** Relative abundance of GTDB-assigned MAG taxa at Guaymas Basin. Abundances are shown (A) for all taxa at the genus level, and (B) for the Archaea at the order level, using read coverage normalized to 100M reads per sample and expressed as a percentage of MAG reads per sample. Relative abundances were averaged for the two samples from the six-day thermocouple array (4561-380 and 4561-384).
