## Additional File 8: Supplementary Discussion for "Global patterns of diversity and metabolism of microbial communities in deep-sea hydrothermal vent deposits"

**Characterization of two new phyla from deep-sea hydrothermal vent deposits**

We identified and characterized two novel bacterial phyla, “JALSQH01” and “JALWCF01” (using GTDB r202 as a reference). These two lineages contained 3 and 13 MAGs, respectively, and formed two distinct phylogenetic clades from other bacterial phyla (both with 100% IQ-TREE UFBoot support). Their relative evolutionary divergence values fell into the phylum range (Table S2) [1]. JALSQH01 MAGs were distributed in MAR, Brothers volcano and ELSC, while JALWCF01 MAGs were only found at ELSC. Metabolic profiling suggested that members of both phyla were associated with aerobic lifestyles, utilizing peptides and polysaccharides as carbon sources (Fig. S6, Table S6). The genomes contained a variety of peptide, amino acid, sugar, polysaccharide, and lipopolysaccharide transporters. Members of both phyla also contained a diverse set of aminotransferases, amino acid utilization pathways to generate acetyl-CoA, and the ability to conduct glycolysis, beta-oxidation, phenol degradation, and benzoyl-CoA reduction suggesting that these organisms can utilize both aliphatic and aromatic fatty acids (Fig. S6, Table S6). Additionally, JALWCF01 MAGs harbored the capacity to oxidize C1 compounds including formate and CO (Fig. S6, Table S6) and fix carbon using the Wood-Ljungdahl pathway. Members of both phyla encoded for cytochrome *bd* ubiquinol oxidase that has high affinity for oxygen, hinting at a microaerobic lifestyle [2]. Beyond central carbon and energy conservation metabolism, members of both phyla contained additional metabolic capacities for hydrogen oxidation, thiosulfate disproportionation, and arsenate, perchlorate, and selenate reduction. The metabolic flexibility of these lineages can potentially provide them with the ability to adapt to ecologically dynamic environments [3, 4].

**Successional changes in deep-sea hydrothermal vent deposits**

The Guaymas samples in this study represent a community succession experiment, as previously described [5, 6]. Samples represented here were collected during the AT15-55 expedition in 2009 using the submersible *Alvin*. A hydrothermal chimney was collected (4559-240), and a thermocouple array placed on the actively venting hydrothermal fluid (Fig. S2). After 6 days, mineral deposits and microbial mats accumulated on the array. Two samples were analyzed from this array (mostly orange mat 4561-380, and orange and white mat 4561-384). The array was placed back on the site, and after 15 days, a small hydrothermal deposit (4571-419) had developed on the array, which was then collected and analyzed.

Although the metagenomes are relatively small, shifts in MAG diversity and abundance over time were evident (Fig. S12A, Table S4). Most notable was an increase in archaeal MAG diversity over time (Fig. S12B). Initial colonization of the array was driven by *Methanocaldococcus* and *Thermococcus*, prevalent Archaea at Guaymas basin vents [7–9]. After 15 days, these taxa were joined by members of Nanoarchaeales, Woesearchaeales, Iainarchaeota and Thermoproteota (Pyrodictiaceae). While the original chimney had examples of all these initial colonizers, it had a much richer diversity of the Thermoproteota (Table S2) and was also colonized by members of the Archaeoglobales and Methanopyrales. Likewise, a greater complexity of Bacteria was seen in the mature chimney, with shifts in key taxa occurring over time. Early colonists included members of the Aquificota and Campylobacterota, which were gradually replaced in abundance by lineages such as the Thermotogota, Desulfobacterota and Caldisericota (Fig. S7, Table S4). Significantly, potentially parasitic Gracilibacteria from the order Absconditabacterales were early colonizers that were subsequently not replaced and were present at all times throughout the colonization experiment. Although generalizations associated with colonization of hydrothermal chimneys are not possible from this one time experiment without replication, it does follow a similar experiment [5], where diversity was measured by 16S rRNA gene fragments (DGGE), and both provided insights into the potential shifts that can occur over time on actively forming chimneys.
